# Scale-dependent feedback between sociality and space use in a long-lived marine predator

**DOI:** 10.64898/2026.02.06.704493

**Authors:** Michaela A. Kratofil, Robin W. Baird, Devin S. Johnson, Colin J. Cornforth, Sabre D. Mahaffy, Michelle Caputo, Jeremy J. Kiszka, Karen K. Martien, Mauricio Cantor

## Abstract

Spatial and social behaviours in animals are intertwined, yet the causal direction of this feedback—and how it varies across scales—remains largely unresolved. Using long-term data from false killer whales (*Pseudorca crassidens*) in the main Hawaiian Islands, we developed and applied a scale-explicit analytical framework to test how social reliance and resource ephemerality govern top-down versus bottom-up processes linking movement and sociality. Movements, association networks, genetic relatedness, and isotopic niches reveal that strong social bonds drive bottom-up emergence of short-term intra-group movements, while ephemeral and likely island-associated prey landscapes impose top-down constraints on inter-group dynamics across scales. These complementary processes generate persistent, fine-scale fidelity within some groups and relatively well-differentiated feeding niches among them. Our findings highlight a general mechanism by which life-history strategies and environmental stochasticity jointly determine the scale and direction of feedback between space use and sociality—shaping population structure and connectivity in mobile social predators.

## 1. Introduction

Animal space use and social behaviour are both shaped by and feed back onto their spatial and social environments (Webber *et al*. 2023). Individual movements depend on the distribution of resources and risks in the spatial environment; this resulting space use determines opportunities for social interactions because individuals can interact if they share the same space (i.e., a top-down effect; Farine 2015; Macdonald & Johnson 2015). At the same time, individuals might incorporate social information indirectly (e.g., eavesdropping on cues) or directly (e.g., tracking signals and cooperating) to find food (Aplin *et al*. 2012), initiate movements (Oestreich *et al*. 2022), and avoid predators (Atkinson *et al*. 2025) or competitors (Mourier *et al*. 2024). Thus, space use inherently emerges from social processes. Inferring the direction of causality in this feedback—whether space use shapes social behaviours or *vice versa*—is critical, given the implications for both immediate effects on survival and long-term consequences for population structuring and persistence (Farine *et al*. 2015; Webber & Vander Wal 2018). Recent conceptualisation of scale-dependence in this spatial-social interface provides a framework for disentangling causal directions, specifically top-down (i.e., coarse scale spatial/social components constrain fine-scale behaviour) versus bottom-up processes (i.e., fine-scale spatial/social components give rise to coarse-scale behaviour; Picardi *et al*. 2024). However, there is limited empirical understanding of what biological or ecological characteristics mediate the predominance of top-down versus bottom-up effects, and how this scale-dependent directionality relates to both short- and long-term consequences.

The degree of dependence on conspecifics throughout an individual’s lifespan (hereafter, “social reliance”) provides a foundation for identifying scale-dependent bottom-up and top-down processes in the spatial-social interface. In several species that forage alone, interactions between individuals result from resource-driven co-occurrence (i.e., predominance of top-down effect of spatial on social behaviour; Carr & Macdonald 1986; Newsome *et al*. 2013), while drivers of interactions in species that live in societies with high fission-fusion dynamics can be context-dependent (e.g., influenced by resource heterogeneity; Peignier *et al*. 2019; Ramos-Fernández *et al*. 2006). In contrast, bottom-up emergence of space use from social processes should be predominant in species that rely on group members to overcome most life-history challenges (Krause & Ruxton 2002). This is especially the case in long-lived and predominantly matrilineal species with advanced cognitive abilities, as longevity enables investment in differentiated social relationships that directly benefit survival (Brent *et al*. 2015; McComb *et al*. 2011; Silk & Hodgson 2021). Inherent in the spectrum of social reliance is resource availability: predictable and defensible resources (e.g., prey) and risks (e.g., predation) can reduce social reliance, whereas heterogenous and ephemeral resource and risk landscapes can increase social reliance (Gaynor *et al*. 2019; Kohles *et al*. 2022; Kun & Dieckmann 2013; Macdonald & Johnson 2015). Thus, on an evolutionary timescale, ecological dynamics have shaped species-level diversity in social complexity (Krause & Ruxton 2002); in contemporary contexts, this established social reliance, combined with variation in the environment, likely mediates the predominance of causal directions in the spatial-social interface.

The synergistic effects of social reliance and resource ephemerality on animal spatial and social behaviour can have implications for long-term population dynamics. Specifically, where long-term variability in spatial (e.g., resource availability) and social environments (e.g., population density) results in unequal fitness among individuals (Bolnick *et al*. 2003), niche variation (e.g., habitat or diet specialisation, site fidelity) can develop at individual- (e.g., Sheppard *et al*. 2018; Strickler 1979) or group-levels where social reliance is strong (e.g., De Stephanis *et al*. 2008; Eguiguren *et al*. 2019). Within-population niche variation can influence the distribution of individuals or groups, and consequently, the probability of inter-individual or inter-group interactions that give rise to population-level social structure (Araújo *et al*. 2011; He *et al*. 2019; Webber *et al*. 2024). This population-level outcome reciprocally influences how individuals navigate their spatial and social environments in the short term (e.g., days to weeks; Cantor *et al*. 2021; Farine *et al*. 2015). Collectively, this implies that causal directions between spatial and social processes could structure a feedback loop linking short-term individual-level patterns with long-term population-level consequences (Figure 1). While similar eco-evolutionary feedback mechanisms have been proposed (Webber & Vander Wal 2018), there has yet to be an explicit examination of how causal directions and scale-dependence regulate this feedback. Identifying the scales at which causal directions are predominant in this feedback would not only shed light on evolutionary causes and consequences in the spatial-social interface but also help elucidate relevant units of conservation and a population’s adaptability to environmental change.

**Figure 1.**
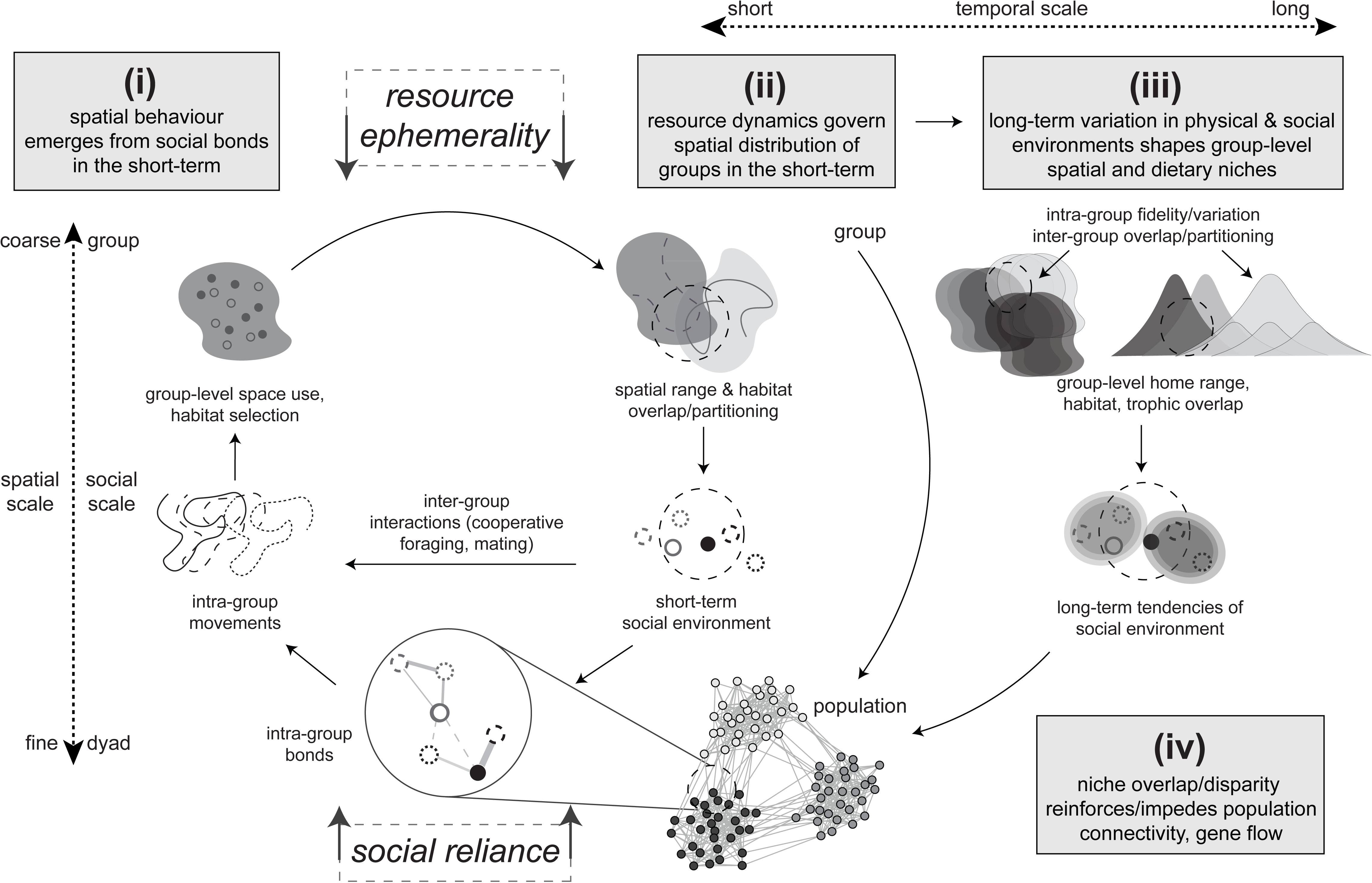
Conceptual diagram proposing social reliance and resource ephemerality as regulators of spatial-social directional feedback mechanisms (i.e., bottom-up versus top-down effects) across spatial, temporal, and social scales. We predict that (i) where social reliance is high, intra-group space use emerges from social bonds in the long-term; (ii) resource ephemerality (interacting with habitat configuration) determines the spatial distribution of social groups in the short-term (e.g., through density-dependent effects), which subsequently influences inter-group interactions in this time scale; (iii) long-term variation in the dynamics of physical and social environments (i.e., predictions (i) and (ii)) shapes niche variation among social groups, which can be measured through intra-group fidelity (or variation) and inter-group overlap in spatial and dietary niches; and (iv) niche overlap/disparity influences long-term tendencies of the social environment (i.e., probability of groups interacting with one another), and thus population connectivity and gene flow.

We empirically explored the scale-dependent feedback between bottom-up and top-down effects on the spatial-social dynamics of a strongly social and long-lived apex predator, the false killer whale *Pseudorca crassidens*. We developed a multi-scale analytical framework combining longitudinal information on social structure, movements, trophic ecology, and genetic relatedness on a small and endangered population occurring around the main Hawaiian Islands. Dyadic social bonds and group membership were obtained from long-term photo-identification (Mahaffy *et al*. 2023) and related to movements inferred from satellite-linked transmitters to derive individual-level spatial behaviours and group-level spatial niches. We used stable carbon and nitrogen bulk isotope analysis from skin biopsy samples to characterize group-level trophic niches and relative dietary contributions of prey. We then used these movement, social, and niche metrics to test our predictions that the longevity and strong social reliance of false killer whales would result in bottom-up emergence of within-group spatial behaviour in the short-term (Figure 1-i), while resource ephemerality and their island-associated habitat configuration would promote/impede between-group interactions in both the short-term (spatial partitioning) (Figure 1-ii) and long-term (niche variation) (Figure 1-iii), with consequences on relatedness and population structure (Figure 1-iv).

## 2. Materials and methods

### 2.1 Study system

False killer whales live into their 60s, are slow to mature, invest significant care into rearing offspring, and are one of a few non-human species that undergo reproductive senescence (Ellis *et al*. 2024; Ferreira *et al*. 2014; Photopoulou *et al*. 2017). This species is resident around the main Hawaiian Islands and have been studied since the late 1990s (Baird *et al*. 2008). The population is small and declining (<160 individuals; Badger *et al*. 2025), and is listed as endangered under the U.S. Endangered Species Act. The population is structured into four stable social groups (hereafter clusters) comprised of kin and non-kin associates (Mahaffy *et al*. 2023). Mating occurs both within and between clusters, and natal fidelity to clusters is strong (Martien *et al*. 2019). While the marine environment around the main Hawaiian Island is generally considered oligotrophic, the island-mass effect promotes enhancement of biological productivity, resulting in more consistent prey availability for marine predators (Gove *et al*. 2016). False killer whales’ prey landscape is, however, much more ephemeral; many of their known prey—such as tunas, billfish, mahimahi *Coryphaena hippurus*—are highly migratory, fast, and difficult to capture. In response, false killer whales undertake extensive movements (up to 150 km/day), cooperatively hunt, and share prey with group members (Baird *et al*. 2008, 2010; Bradford *et al*. 2014).

### 2.2 Empirical data

#### 2.2.1 Satellite tagging and tissue sampling

Empirical data types and processing workflows are summarized in Figure S1. Satellite tagging and biopsy sampling were undertaken during small boat-based surveys in Hawaiʻi from 2007 to 2025 (Baird *et al*. 2013b, 2024). Tagging (Baird *et al*. 2010, 2012), biopsy sampling (Ylitalo *et al*. 2009), stable isotope (Kratofil *et al*. 2020; Supporting Information), and genetic (Martien *et al*. 2014, 2019) laboratory procedures followed standard protocols.

#### 2.2.2 Deriving individual-level traits

Photographs of all tagged and biopsy sampled individuals were matched to a long-term photo-identification catalogue to determine individual identity and social cluster membership (Baird *et al*. 2008; Mahaffy *et al*. 2023). We used a temporally-constrained half-weight index (HWI) as a metric for dyadic associations, which accounts for periods when individuals’ sighting histories overlapped (Whitehead 2009). Sex was either determined genetically from biopsies, inferred from morphological (e.g., head shape) or social characteristics (i.e., individuals consistently accompanied by calves), or assigned as “unknown”. Age class was determined from photo-identification-derived metrics (see Kratofil *et al*. 2026). Movement was tracked using satellite-linked transmitters (n=69 individuals; median transmission time=41 days, range=9-229; Cluster 1, n=31 deployments/25 individuals; Cluster 2, n=9/9; Cluster 3, n=20/20; Cluster 4, n=9/8). The bulk stable carbon and nitrogen isotopes dataset included 80 samples (Cluster 1, n=27 samples/22 individuals; Cluster 2, n=22/22; Cluster 3, n=12/10; Cluster 4, n=19/19). Pairwise genetic relatedness was available for a subset of tagged individuals (n=23) and calculated using the data and methods from Martien *et al*. (2014, 2019).

### 2.3 Quantifying individual-level spatial metrics

We derived individual-level spatial metrics (movements, ranges, habitat selection) from satellite tracks as the basis for subsequent analyses (Figure S1). All analyses were completed in R v4.5.0 (R Core Team 2025). Tag location data were processed following Kratofil et al. (2023) and subsequently fit to continuous-time movement models using the *ctmm* and *ctmmUtils* packages (Fleming & Calabrese 2023; Johnson & London 2025; Supporting Information). All models described in the following sections follow the same fitting (see Supporting Information) and performance check procedures (e.g., R-hat values<1.05, mixing of chains, and posterior predictive checks) unless otherwise specified.

We characterized the spatial ranges used by tagged individuals through optimally weighted and area-corrected autocorrelated kernel density estimators (wAKDE UDs) in the *ctmm* package (Fleming & Calabrese 2023), which accounts for irregular sampling in time (Fleming *et al*. 2018; Silva *et al*. 2022). Because these tracks only reflect a comparatively brief portion of an individual’s life, we refrain from referring to these ranges as “home ranges”, which reflect long-term space use (Péron 2019). We defined core ranges as the 50% isopleth of the wAKDE UD.

We quantified individual-level habitat selection through a “used-available” framework within each tagged individual’s spatial range (i.e., third order, Manly *et al*. 2002), which we defined as the 95% isopleth of their wAKDE UD. Tag locations represented “used” locations, and “available” locations were randomly sampled within each individual’s spatial range at a 1:20 ratio (Northrup *et al*. 2013). We used 6-hourly locations (predicted using *ctmm*) to mitigate autocorrelation effects while also reflecting a relevant scale of habitat selection given false killer whale movement rates. Locations with error ellipses greater than 20 km were removed prior to modelling. Environmental variables related to biological productivity were extracted for each location; where two variables were correlated (Pearson’s correlation>0.5), only one was included for subsequent modelling, and selected based on the most direct ecological interpretation (Table S1). Resource selection functions (RSFs) were fit as logistic regressions (Bernoulli distribution with logit link; Table S2; Manly *et al*. 2002) in the *brms* package (Bürkner 2018).

### 2.4 Short-term within-cluster emergence and between-cluster overlap of space use

To test predictions (i) and (ii) (Figure 1), we compared spatial metrics of individuals tracked simultaneously in relation to their pairwise HWIs and cluster membership. We first calculated the pairwise distances between temporally overlapping tracks using the *ctmm* package (Fleming & Calabrese 2023). We then modelled pairwise median distance apart in relation to HWI using a Bayesian generalized linear mixed effects model (bGLMM) in *brms* (Bürkner 2018) with a gamma response distribution (log link) and multi-membership random effect for the pair of catalogued identifications (Hart *et al*. 2022; Table S2). We limited the dataset to individuals sighted five or more times.

We then selected seven periods with reasonable coverage of tagged individuals (1-4 months) within the same cluster or of different clusters to examine all spatial metrics. For these cases, we truncated tracks so that all tracks covered the exact same period; all spatial metrics were re-calculated with these equal-length tracks and qualitatively assessed (Figure S1). Overlap between spatial ranges was estimated using the bias-corrected Bhattacharyya’s coefficient (BC) within the *ctmm* package (Fleming & Calabrese 2023; Winner *et al*. 2018). To estimate overlap between core ranges, we first calculated the area of intersection between core range polygons using the *sf* package (Pebesma 2018), and then defined overlap as two times the ratio of this area out of the combined core range areas for each pair.

### 2.5 Long-term within-cluster fidelity and between-cluster overlap

For prediction (iii) (Figure 1), we quantified long-term spatial and core range overlap (described above) between all pairs of tagged individuals across the entire study period. We fit bGLMMs to model spatial range overlap (BC; beta distribution, identity link function) in relation to unique cluster pairs (e.g., C1-C1, C1-C2, etc.) and a multi-membership random effect for each pair of individuals. We also included fixed effects for year (same/different) and difference in transmission duration (scaled) to account for any effects these variables may have on our inferences. We ran separate models on a subset of dyads for which pairwise genetic relatedness information was available to assess the potential influence of genetic similarity on space use (Table S2). We excluded pairs where the mean overlap estimates suggested range overlap (BC >0.5) but there was low confidence in this estimate (lower CI <0.1; Winner *et al*. 2018). The same model was fit for core range overlap, albeit with a zero-inflated beta distribution (Table S2).

We additionally assessed spatial fidelity by comparing spatial metrics within six individuals that have been tagged more than once during the study period. Within each individual, we truncated their unique tracks to the length of the shortest track and recalculated spatial metrics (following section 2.4).

### 2.6 Long-term cluster-level spatial and dietary niches

#### 2.6.1 Spatial niches

We derived cluster-level home ranges from individual-level wAKDE UDs using the *ctmm* package (Fleming & Calabrese 2023) and considered the 50% isopleth of these outputs as the cluster-level core range. Cluster-level home and core range overlap were determined following the same overlap methods described in section 2.4.

Cluster-level habitat selection was estimated using a two-stage approach following Johnson *et al*. (2022). The individual-level RSFs from section 2.3 constituted the first stage; posterior means and covariance matrices were extracted from the posterior samples and subsequently modelled as observations in a meta-mixed effects regression model (with a multivariate normal distribution) in relation to cluster membership and a random effect for tag ID; models were fit using the *mixmeta* package (Sera *et al*. 2019).

#### 2.6.2 Isotopic niches and diet estimates

No temporal or demographic effects on *δ*^13^C and *δ*^15^N values were found (Supporting Information). We estimated bivariate (*δ*^13^C and *δ*^15^N) isotopic niche space (95% probability overall region) and pairwise niche overlap (95% and 40% probability region) among social clusters using the *nicheROVER* R package (Lysy *et al*. 2023; Swanson *et al*. 2015).

We used stable isotope mixing models to estimate the relative dietary proportions of prey among social clusters. *δ*^13^C and *δ*^15^N values of 13 known false killer whale prey were compiled from the literature (Table S6) and aggregated into groups *a posteriori* to increase the discrimination power and interpretability of the mixing model (Phillips *et al*. 2005, 2014; Stock *et al*. 2018). We defined four groups based largely on ecological and functional traits (Table S7): (1) epipelagic predatory fish; (2) mesopelagic predatory fish; (3) reef-associated predatory fish, and (4) mesopelagic cephalopods. Mixing models were fit using the *MixSIAR* package (Stock *et al*. 2018; Stock & Semmens 2016), with social cluster used as a random effect. We ran models with 300,000 iterations (200,000 burn-in, thinned every 50 samples) over three Markov chain Monte Carlo (MCMC) chains. Model convergence was checked using the R-hat (estimates <1.05) and Geweke diagnostic values (<5% of estimates less than ± 1.96).

### 2.7 Population connectivity

To assess prediction (iv), we derived the conditional distribution of encounters (CDE) across all individuals, which identifies areas with the highest probability of individuals encountering one another (Fleming & Calabrese 2023; Noonan *et al*. 2021). We then mapped cluster-level core ranges on top of the population-wide CDE and descriptively related CDE hotspots to core range overlaps and the composition of mitochondrial haplotypes among clusters (Mahaffy *et al*. 2023).

## 3 Results

### 3.1 Short-term space use is an emergent property of social bonds

Pairs of tagged false killer whales (n=80 pairs with ≥10 d overlap) that were more strongly associated were more spatially cohesive (posterior median: −4.51, 95% credible interval (CrI): [− 5.64; −3.36]; Figure 2, Table S3). There were some pairs who were associates (HWI >0.2) in the same cluster (either Cluster 1 or 3) but did not remain spatially associated throughout their common tracking period (Figure S2). A single pair involving individuals from different clusters (Clusters 1-3) had a median distance of less than 10 km over 10 days. Pairs from Clusters 2 and 4 exhibited cohesive movements more than that observed for Clusters 1 and 3 (Figures S2, S3). Even when same-cluster pairs were not consistently spatially associated, spatial fidelity persisted (Figure 3a,b), and RSF coefficients were highly similar (Figure 3c,d, S3).

**Figure 2.**
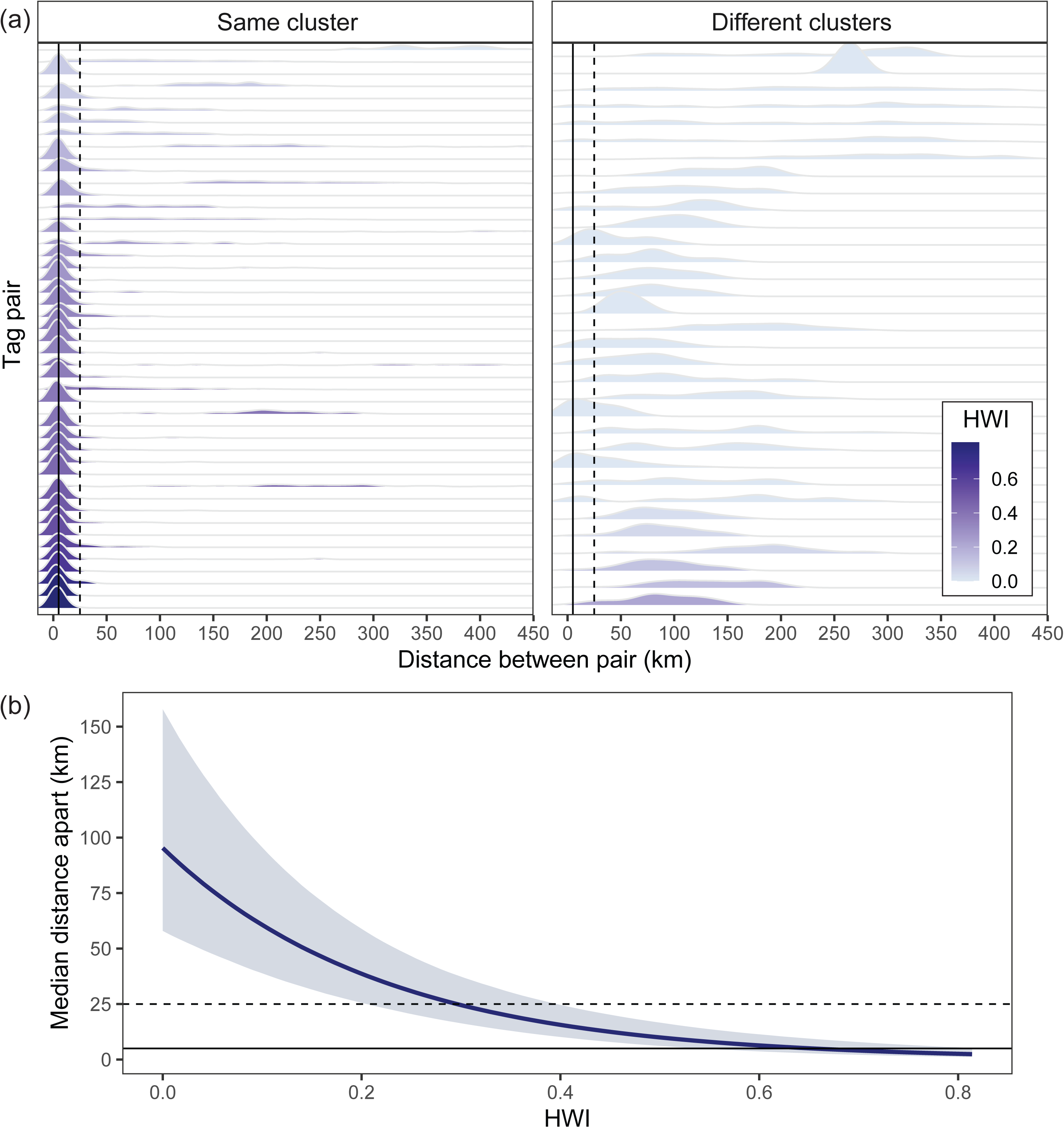
Relationship between social associations and spatial cohesion in tagged false killer whales. (a) Distribution of distance between tagged dyads sorted by decreasing pairwise association strength (half-weight index, HWI) for same-cluster pairs (left) and different-cluster pairs (right; vertical solid line = 5 km; vertical dashed line = 25 km); (b) conditional effects plot from the fitted model relating median distance between dyads and pairwise association strength (dark line = estimate; shaded ribbon = 95% credible interval; solid lines = 5 km; dashed lines = 25 km).

**Figure 3.**
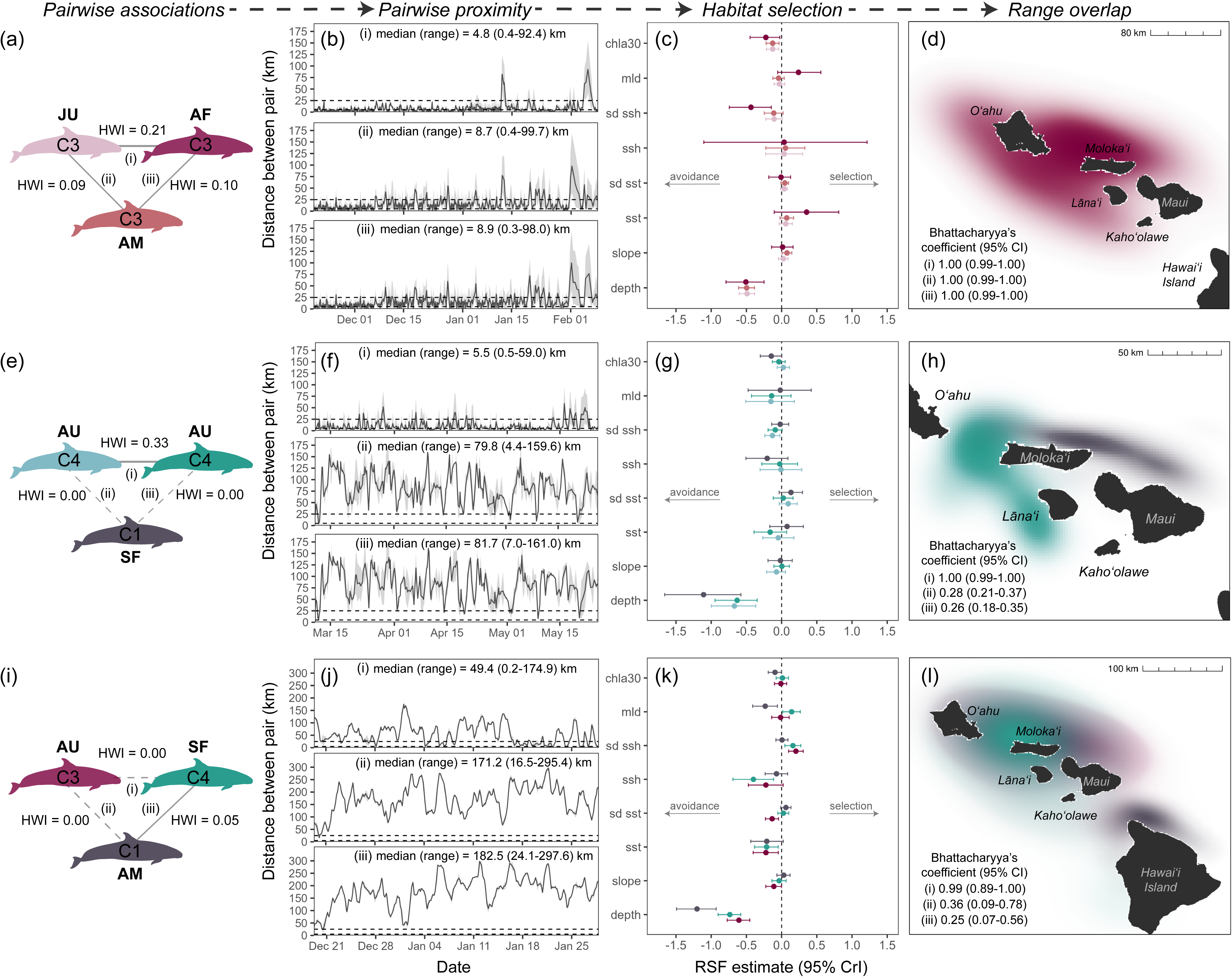
Short-term intra- and inter-cluster false killer whale social associations and spatial behaviour across scales. Emergence of intra-cluster space use from social bonds (a-h) from fine to coarse scales (left to right) and between-cluster partitioning (e-l) in the short-term. Each panel shows (1^st^ column) pairwise association strength (half-weight index, HWI) among the focal individuals (i, ii, iii) with cluster membership (colour and “C#”) and age-sex class (A = adult, S = sub-adult, J = juvenile; F = female, M = male, U = unknown); (2^nd^ column) pairwise proximity (distance between each pair) for the period of overlap, with 5-km and 25-km indicated by horizontal dashed lines; (3^rd^ column) habitat selection coefficients of individuals (estimates and 95% credible intervals, CrI; mld = mixed layer depth, chla30 = 30-d lagged surface chlorophyll-a concentration, sd ssh = standard deviation of sea surface height, ssh = sea surface height, sd sst = standard deviation of sea surface temperature, sst = sea surface temperature, slope = seafloor slope, depth = seafloor depth); and (4^th^ column) spatial range overlap, with the Bhattacharyya’s coefficient values and 95% confidence intervals (CI) shown for each pair.

### 3.2 Short-term spatial partitioning between social clusters

Spatial overlap between clusters was typically low during restricted periods (Figure 3e-l, S3). Where overlap occurred, individuals of different clusters rarely, if ever, encountered one another. For example, individuals from Cluster 3 and Cluster 4 had high range overlap, but only occasionally came within 5 km of one another (Figure 3i-l). RSF coefficients between clusters during restricted periods were similar; while variation in individual coefficient estimates occurred, the 95% CrIs between individuals were highly overlapping (Figure 3e-l, S3).

### 3.3 High short-term spatial and dietary overlap but distinct long-term core niches

Long-term within-cluster fidelity and between-cluster spatial overlap were high (Figure 4a). Unique cluster pairs had the strongest effect on spatial range overlap, specifically pairs from Cluster 4 (Figure 4a; Table S3). Cluster 4 also exhibited the lowest posterior mean, but the largest posterior uncertainty in range fidelity (Figure 4a). Core range fidelity and overlap were lower than that for spatial ranges (Figure 4a). Within-cluster fidelity was highest for Clusters 2 and 4 (albeit with overlapping credible intervals), and between-cluster overlap was lowest between Cluster pairs 1-4, 2-4, and 3-4 (Figure 4a). For both models, there was either no (core) or minimal (spatial) estimated effect of tag duration difference on overlap, while overlap was higher between pairs tagged in the same year (Table S3). There was no evidence for an effect of genetic relatedness on spatial nor core range overlap for relevant pairs (Table S3).

**Figure 4.**
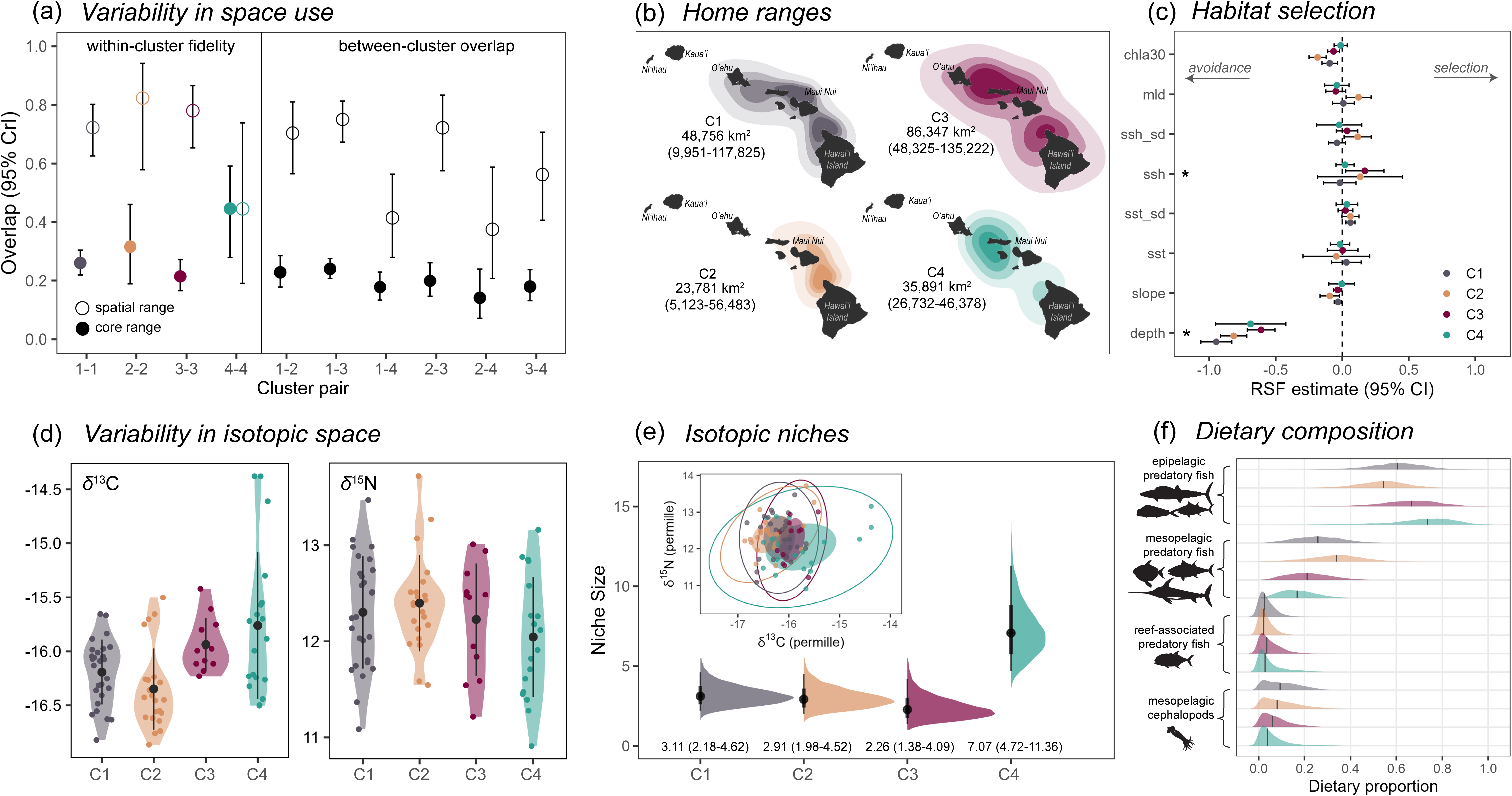
Spatial and trophic niches among false killer whale social clusters. (a) Conditional effects (posterior estimates, 95% credible intervals, CrI) of pairs of social clusters (x-axis) on spatial (open circles) and core range (filled circles) overlap models, showing within-cluster fidelity and between-cluster overlap; (b) cluster-level home ranges defined by population-level wAKDEs (95%, 75%, 50%, and 25% contours shown), with area estimates and associated 95% confidence intervals (CI) shown for each cluster. Maui Nui includes the islands of Maui, Molokaʻi, Lānaʻi, and Kahoʻolawe (see Figure 3); (c) cluster-level habitat selection coefficients (estimate, 95% confidence interval, CI; mld = mixed layer depth, chla30 = 30-d lagged surface chlorophyll-a concentration, sd ssh = standard deviation of sea surface height, ssh = sea surface height, sd sst = standard deviation of sea surface temperature, sst = sea surface temperature, slope = seafloor slope, depth = seafloor depth) with variables where cluster membership had a significant effect on selection (relative to Cluster 1, from mixed-meta regression) indicated by asterisks; (d) distribution of *δ*^13^C and *δ*^15^N isotopes by social cluster (mean and standard deviation plotted in centre); (e) cluster-level posterior isotopic niche size (point = median; line range = 95% CrI; shown in text) and inset ellipses (overall/95% outlined; core/40% shaded); (f) posterior densities of dietary proportions of prey groups among social clusters (median shown by vertical line) based on stable isotopes analysis of skin samples.

Cluster-level home ranges were variable in size and overlap (Figure 4a,b). Clusters 1 and 3 had the largest home ranges and Clusters 2 and 4 had the smallest (Figure 4b). There were minor differences in RSF coefficients between clusters, however, confidence intervals almost always overlapped (Figure 4c). Selection for shallower waters dominated habitat selection patterns (*depth*: Figure 4c), while selection for dynamic habitat features was minimal at this scale (Figure 4c). Cluster-level isotopic niches highly overlapped among all clusters (Figure 4d; Table S5). Isotopic niche spaces were similar in size among Clusters 1, 2 and 3, whereas Cluster 4 had a wider isotopic niche, mainly driven by a broader range in *δ*^13^C values (Figure 4e). Estimated prey contributions were similar among clusters—leading with epipelagic predatory fish, then mesopelagic predatory fish, and small proportions (<20%) of reef-associated predatory fish and mesopelagic cephalopods (Figure 4f; Table S8). Clusters 1 and 2 had slightly higher proportions of mesopelagic predatory fish compared to the other clusters, while Clusters 3 and 4 had slightly higher proportions of epipelagic predatory fish (Figure 4f). These results were robust to prey grouping choices and source data (Supporting Information).

Within-individual spatial range fidelity was variable. Only three of six individuals had high overlap between successive spatial ranges (HIPc145, HIPc202, HIPc272; Figure S4), one of which was tagged three times (HIPc145) and only the third range was more disparate (Figure S4).

### 3.4 Range-wide hotspots and habitat structure mediate between-cluster dynamics

The CDE revealed two primary areas where false killer whales are most likely to co-occur: one off northwestern Hawaiʻi Island (including the ʻAlenuihāhā Channel) and the other being the Kaiwi Channel, between Oʻahu and Molokaʻi, both of which are characterized by high wind stress and current velocity (Figure 5). The core ranges of Cluster 1, 2, and 3 overlapped with the region off Hawaiʻi Island, while those of Cluster 1, 3, and 4 included the Kaiwi Channel hotspot (Figure 5). Clusters 3 and 4 have only mitochondrial haplotype 1, while Clusters 1 and 2 have both haplotype 1 and 2 (Mahaffy *et al*. 2023).

**Figure 5.**
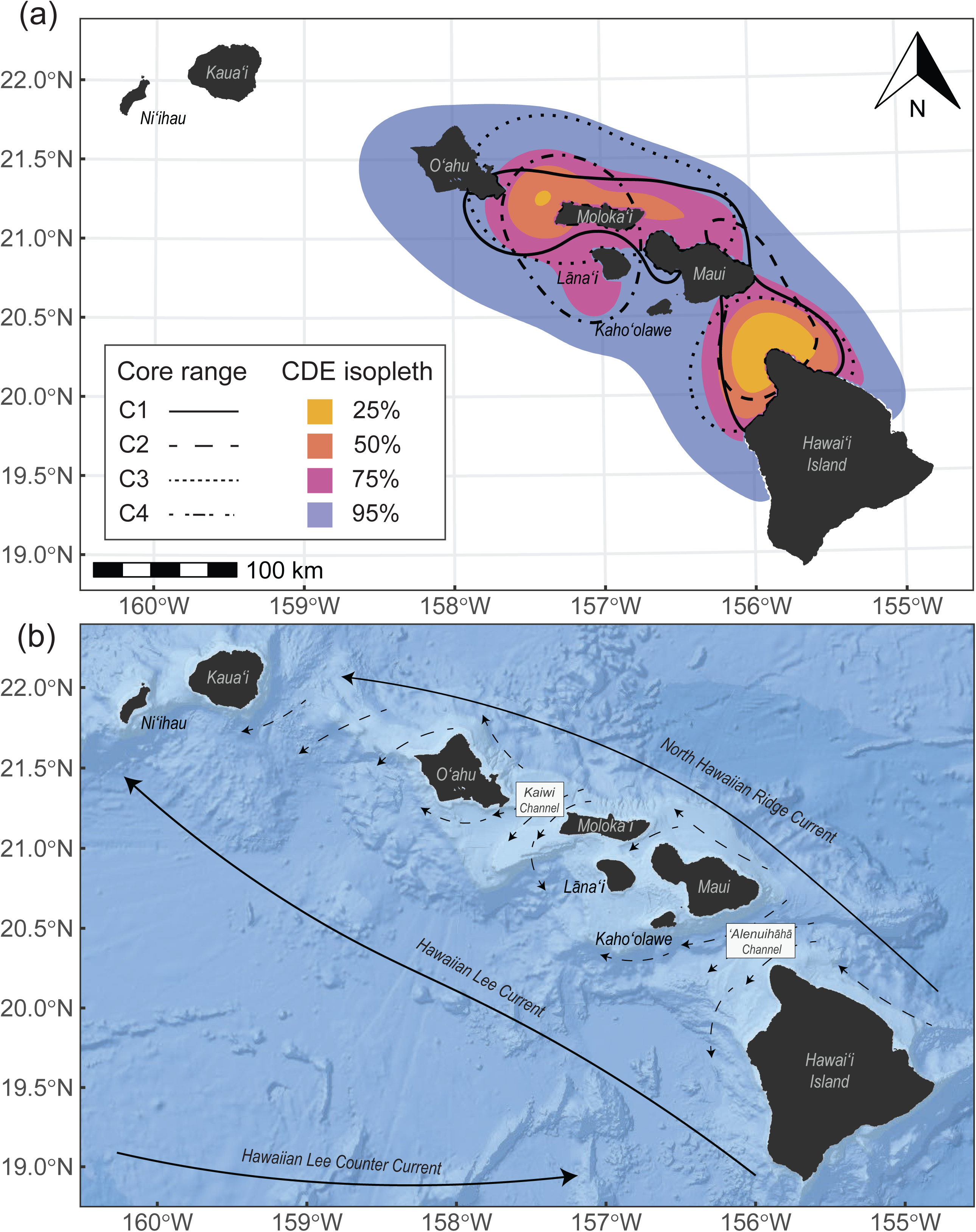
Co-occurrence hotspots of false killer whales in relation to broad scale habitat features around the main Hawaiian Islands. (a) Overlap between population-wide spatial hotspots (proxied by the conditional distribution of encounters, CDE) and social cluster-level core ranges (C1 = Cluster 1, C2 = Cluster 2, C3 = Cluster 3, C4 = Cluster 4; outlined). (b) Primary currents (solid lines) and wind, current directions (dashed lines), and channels (outlined boxes) relevant to observed space use in panel (a). Current and wind lines were adapted from Lindo-Atichati *et al*. (2020). Basemap image in (b) is the intellectual property of Esri and is used herein with permission (Copyright 2025 Esri and its licensors, all rights reserved).

## 4 Discussion

Using a novel scale-explicit conceptual framework with a longitudinal study of false killer whales, we demonstrated how strong sociality yields bottom-up emergence of intra-social cluster spatial behaviour from social bonds in the short term, while their ephemeral resource landscape regulates both short- and long-term inter-cluster dynamics. Most importantly, we show how these bottom-up and top-down processes may synergistically affect long-term population structure.

### 4.1 Social reliance mediates bottom-up emergence of spatial behaviour

As we hypothesised, false killer whale dyads that have stronger associations tended to move more cohesively. In highly social animals, movements reflect collective decisions and coordination among group members (Strandburg-Peshkin *et al*. 2015), a process that requires some level of spatial cohesion (Conradt & Roper 2005). Spatial cohesion is especially important for predators that cooperatively hunt, share prey, or care for offspring (Hansen *et al*. 2023), as effective group coordination underlies individual success and survival. Consequently, cohesive movements should arise from the social bonds that form from the reliance on group members for these behaviours, and our findings are consistent with this expected directionality. While spatial cohesion observed herein was often over several kilometres, false killer whales likely acoustically communicate over this range, as they are known to exhibit coordinated movements over extensive distances (>20 km; Baird *et al*. 2008; Bradford *et al*. 2014).

The dyads that did not conform to this relationship (a subset of Cluster 1 and 3 members) could reflect cluster-specific behavioural properties unmeasured in our study. For example, group size or composition can influence intra-group spatial behaviour if sub-optimal sizes or certain demographic classes (e.g., younger, slower individuals) decrease the efficiency or performance of group-level decision making (Cantor *et al*. 2020; Fryxell *et al*. 2025; Papageorgiou & Farine 2020) or are more likely to disperse (e.g., males; Clutton-Brock 2016). Evidence for sub-clustering within Cluster 3 (Mahaffy *et al*. 2023) may explain some dyad’s more diffuse spatial association patterns, while higher association strengths within Clusters 2 and 4 (Mahaffy *et al*. 2023) could explain these groups’ more cohesive movements. Given that mating occurs both within and between clusters (Martien *et al*. 2019), sex and reproductive status may play a role in dyad-level movement patterns, as some less spatially cohesive dyads that occasionally associated in space included an adult male (Figure 3). While social processes shape fine-scale cohesion within clusters, the broader structuring of space among clusters is likely governed by ecological constraints, particularly the ephemerality and distribution of resources.

### 4.2 Resource ephemerality regulates short-term inter-cluster space use

Social clusters spatially partitioned themselves in the short-term at different scales. Large-scale spatial partitioning (e.g., low range overlap over weeks to months) theoretically reduces indirect intra-cluster resource competition (Fretwell & Lucas 1969), particularly when resources are ephemeral and distributed across disparate patches of sufficient quality to sustain separate foraging groups. Indeed, our findings from stable isotopes show that all false killer whale clusters feed across the same prey trophic levels, suggesting that large-scale spatial partitioning may be a necessary tactic to maintain intra-cluster foraging success when prey is patchily distributed and relatively unpredictable.

In contrast, cases where different clusters exhibited high spatial range overlap but tagged dyads were rarely in close proximity (i.e., small-scale partitioning) could reflect temporary access to high-quality prey patches that can sustain multiple clusters simultaneously (Macdonald 1983). Such ephemeral resource hotspots may promote temporary inter-cluster associations (Papageorgiou *et al*. 2019). Because false killer whales are highly social but not territorial, such periods of high spatial overlap could also provide opportunities for mutually beneficial inter-cluster interactions, such as cooperative foraging, social learning, and mating. In this sense, foraging hotspots may function not only as ecological resources but also as social arenas, a dual role increasingly recognized in highly social terrestrial mammals (Strauss *et al*. 2024).

While logistically infeasible in our system, measuring patch quality would provide a more robust basis for testing resource ephemerality and density-dependent effects on false killer whale inter-cluster dynamics. Nonetheless, high similarity in habitat selection across clusters during short periods—combined with the isotopic evidence—indicate that clusters share the same prey base. These results collectively support the hypothesis that resource ephemerality exerts a top-down influence on inter-cluster spatial and social behaviours in the short-term (Figure 1-ii). Over longer timescales, however, resource dynamics may not only modulate short-term associations but also shape persistent patterns of spatial fidelity and niche differentiation among clusters.

### 4.3 Long-term integration of spatial, social, and ecological processes

Consistent use of space over time can provide a variety of benefits, including efficient resource exploitation, familiarity with resource and predator landscapes, and reduced movement costs (Piper 2011; Switzer 1993). Over the 18-year study period, within-cluster spatial fidelity was most variable in Clusters 1 and 3 and least variable in Clusters 2 and 4. With the exception of Cluster 4, intra-cluster fidelity was highly comparable to inter-cluster overlap, suggesting that, in the long term, the population shares a common geographic range, while Cluster 4 occupies a more distinct spatial niche. Lower core range fidelity overall indicates that a cluster’s core area at a given time may be more variable and sensitive to fluctuations in short-term prey availability and density-dependent effects, with some clusters (2, 4) exhibiting greater temporal consistency than others (although see discussion on caveats below). The low within-individual spatial fidelity among some re-tagged whales, and lack of effect of genetic similarity on range overlap, further supports the notion that short-term space use can shift dynamically in response to local conditions.

Similar to the short-term patterns, these long-term differences in space use and fidelity among clusters likely reflect both ecological variability and cluster-specific behavioural properties. For example, within-group core range fidelity is greater when group composition is stable in terrestrial group-living species (Ogino *et al*. 2025), suggesting that demographic turnovers can influence how groups make collective decisions. In species that maintain stable social membership, group turnover can imply the loss of older, experienced individuals with knowledge of alternative resource hotspots (Kopf *et al*. 2024), further influencing collective movement and cluster-level space use; this is a plausible process for our study population, which has declined over the past decade (Badger *et al*. 2025). Thus, both social and environmental processes that occur at finer temporal scales—such as group composition and resource variability—may contribute to the observed temporal variability in cluster-level space use, while enduring cluster-level behavioural traits may underpin the long-term temporal consistency seen in Clusters 2 and 4.

At coarser scales, cluster-level home and core ranges mirrored these spatial patterns. Whales from Clusters 1 and 3 occupied larger, more overlapping areas, whereas the ranges of those from Clusters 2 and 4 were more restricted. All clusters selected for similar habitats and had largely similar trophic ecologies, with some differences in habitat variables (primarily depth), isotopic niche widths, and prey groups consumed. While overlapping cluster-level niches are expected given that high prey field stochasticity constrains resource specialisation in marine predators (Courbin *et al*. 2018), we further found that Clusters 2 and 4 appear to have adopted more distinct spatial and dietary niches than Clusters 1 and 3. Cluster 4 exhibits the widest isotopic niche—driven by a broader range of *δ*^13^C values consistent with feeding on more nearshore prey. This aligns with Cluster 4’s home and core ranges encompassing shallower, coastal habitats than the other clusters. These findings, combined with limited evidence for use of other islands outside of their core range, potentially suggest that increased site fidelity and diversified trophic niche may represent effective strategies for optimizing spatial and social environments.

While limited tag sample sizes for Cluster 2 and Cluster 4 (9 tracks each) introduce caveats, both clusters included very long deployments (>80 days) showing consistent ranging behaviour across years. These patterns, coupled with rare sightings of these clusters outside their core ranges (Mahaffy *et al*. 2023), reinforce the hypothesis that spatial and dietary differentiation among clusters is stable over time. Future high-resolution tracking could test whether fine-scale habitat preferences further partition space among clusters within their shared ranges. Collectively, the current results suggest that both short- and long-term processes shape how social clusters interact in space, ultimately influencing the degree of connectivity and potential for population structuring.

### 4.4 Linking spatial-social dynamics to population structuring and connectivity

How individuals and groups navigate their spatial and social environments—with both short-term decisions and in long-term strategies—determines the probability of encounters among individuals or groups (Aureli *et al*. 2008; Webber *et al*. 2023). These processes collectively shape population-level social structure, including patterns of information transfer, mating and ultimately gene flow (Albery *et al*. 2021; He *et al*. 2019; Spiegel *et al*. 2017). In Hawaiian false killer whales, overlap between cluster-level core ranges was highest in two population-wide hotspots (proxied by the CDE), suggesting that these areas are consistently valuable and may serve as arenas for inter-cluster encounters. Critically, pairwise core range overlaps correspond with cluster-level mitochondrial haplotype composition (Mahaffy *et al*. 2023). While these are broad descriptive correlations, they offer rare empirical evidence that feedback mechanisms between spatial and social processes across scales can shape both population connectivity and genetic structuring.

The unique habitat configuration of the Hawaiian Islands likely reinforces these patterns. Compared to the surrounding open-ocean, the archipelago likely promotes site fidelity through stable local resource patches and memory-based foraging, while the distances separating them are sufficient to limit constant inter-cluster contact—allowing for partial divergence among social clusters that can persist within distinct portions of the island chain. At the same time, proximity among island areas (particularly Oʻahu, Maui Nui, Hawaiʻi) maintains opportunities for social contact, information sharing, and mating, thereby at least partially mitigating potential inbreeding risks. Interestingly, while false killer whales use all the main Hawaiian Islands, their home ranges remain concentrated in these central islands, possibly reflecting higher resource abundance or the shorter inter-island distances. Alternatively, limited ranging near Kauaʻi and Niʻihau may instead result from competition with the Northwestern Hawaiian Islands false killer whale population (Baird *et al*. 2013a; Kratofil *et al*. 2023) or other ecological differences yet to be examined. Collectively, these results indicate that spatial and social processes interact across scales to generate a balance between cohesion and differentiation within this population—a dynamic equilibrium that promotes both local adaptation and population connectivity.

### 4.5 Conclusions

Together, our findings illuminate general mechanisms by which life-history strategies and environmental stochasticity jointly determine both the scale and direction of feedback links between space use and sociality. In Hawaiian false killer whales, this interplay yields a dynamic balance between cohesion and differentiation—promoting population connectivity while maintaining socially mediated structure. Beyond this system, this framework offers a tractable approach for examining how behaviour scales up to shape population structure in mobile species. Identifying the relative influence of these feedback mechanisms across systems of varying social complexity, resource predictability, and habitat heterogeneity will be key to revealing more general principles linking individual behaviour, group dynamics, and the evolution of population structure.

## Acknowledgements

We are grateful to Daniel Webster, Allan Ligon, and Greg Schorr for deploying satellite tags and collecting biopsy samples earlier in the study period. We thank Daniel Palacios, Tiffany Garcia, Will White, Labirinto, Damien Farine & the Farine Lab, Clarissa Teixeira, and Josh Stewart for providing feedback on analyses, and Clara Bird and Janelle Badger for reviewing the draft manuscript. We appreciate 2008-2011 false killer whale stable isotope data provided by Gina Ylitalo, Doug Burrows, and Jonelle Gates of the Northwest Fisheries Science Center, Aaron Carlisle and Brian Popp for providing prey isotope data, and PIFSC for providing tag data for one deployment. MAK was supported by National Science Foundation Graduate Research Fellowship Program Award #1840998, the OSU Marine Mammal Institute Gray Whale License Plate Program, the ARCS Oregon Foundation (Mike & Sheila Goodwin), and PIFSC. Training in Bayesian modelling supported by the NSF award #2042028 enhanced this work. Fieldwork where tagging, sampling, and photo-identification operations were undertaken was supported by the U.S. Navy (Pacific Fleet, Living Marine Resources, Office of Naval Research), NOAA Fisheries (PIFSC, SWFSC, Bycatch Reduction Engineering Program), Dolphin Quest, and the State of Hawaiʻi. MCantor was supported, in part, by the Marine Mammal Institute and College of Agricultural Sciences at Oregon State University, and by the Oregon Agricultural Experiment Station with funding from the Hatch Act capacity funding program (NI25HFPXXXXXG022, NI25HMFPXXXXG029) from the USDA National Institute of Food and Agriculture.

## Conflict of Interest

The authors declare no conflicts of interest.

## Notes

### Competing Interest Statement

The authors have declared no competing interest.

