## Supporting Information for "Scale-dependent feedback between sociality and space use in a long-lived marine predator"

This file includes:

- Supporting methods and results
- Figures S1-S7
- Tables S1-S10
- Supporting information references

Supporting Information S1: Analytical framework

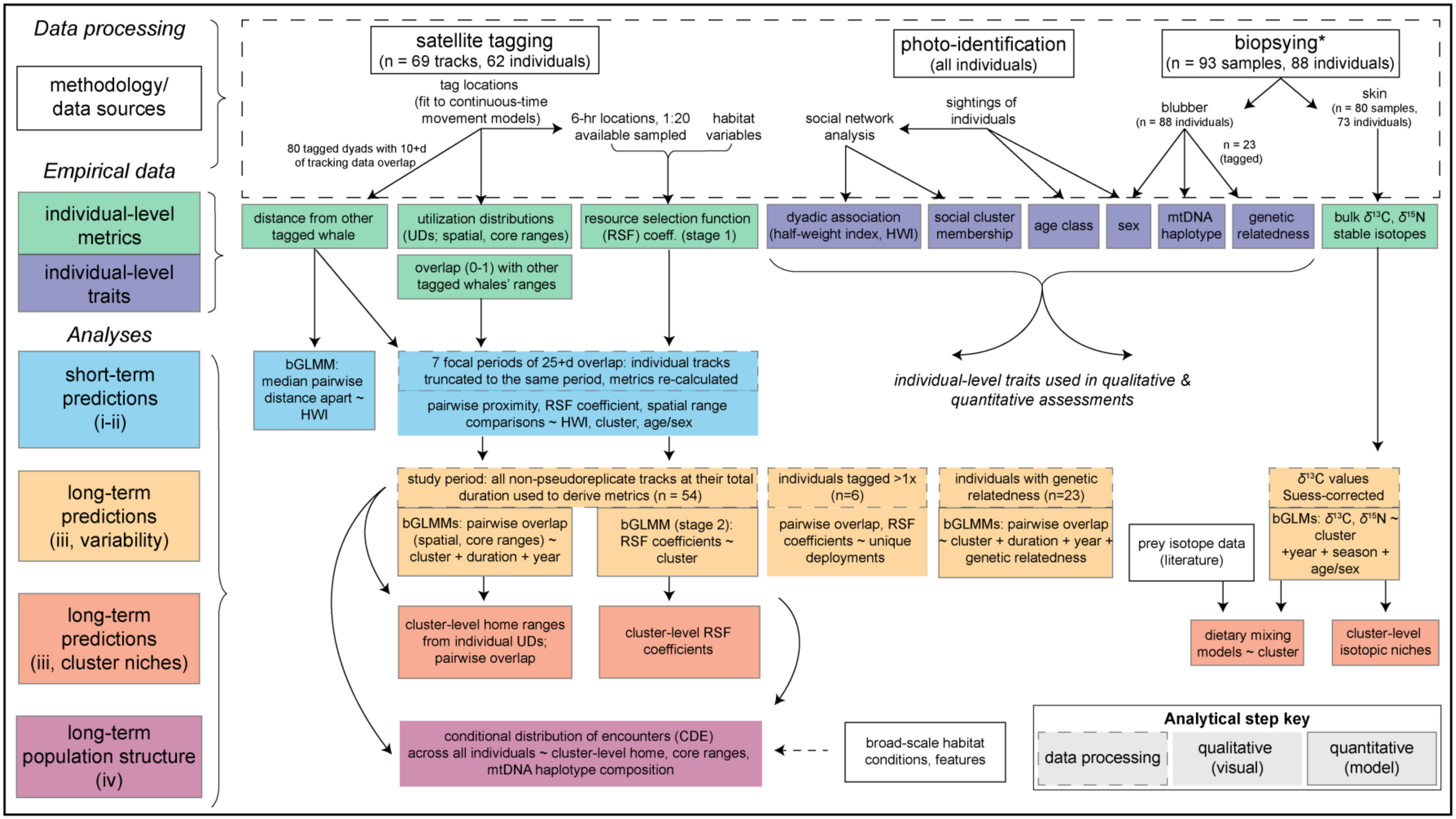


**Figure S1.** Conceptual overview of data sources and processing, resulting individual-level empirical data, and subsequent analytical steps used to test predictions corresponding to Figure 1 in the main text. *All biopsied individuals have genetic sex and mtDNA haplotype. Genetic pairwise relatedness was only assessed for tagged whale space use, and thus the sample size reflects the number of whales tracked, biopsied, and for which pairwise genetic relatedness information is available. The stable isotopes dataset primarily reflects data from individuals not tagged (n=56); while some individuals with stable isotopes were also tagged (n=17), the temporal window reflected by the two data sources does not overlap. Definitions: mtDNA = mitochondrial DNA; bGLMM = Bayesian Generalised Linear Mixed Effects Model; bGLM = Bayesian Generalised Linear Model; spatial range overlap = estimated through Bhattacharyya's coefficient; core range overlap = estimated as twice the area of intersection divided by the combined area of each dyad's core ranges.

Supporting Information S2: Biopsy sample collection and laboratory procedures

*Stable isotope analysis*

Biopsy and skin sample preparation methods follow that reported in Ylitalo et al. (2009) and Kratofil et al. (2020). Skin samples from Kratofil et al. (2020) (n = 54; 2008-2011) were included in this study as well as 26 additional samples collected between 2015 and 2020. The Kratofil et al. (2020) samples were analysed for bulk carbon and nitrogen ($\delta^{13}C$ and $\delta^{15}N$) stable isotopes at NOAA Northwest Fisheries Science Center and the more recent 26 samples at the Institute of Environment's Stable Isotope Facility at Florida International University (North Miami, FL). Stable isotope analyses of skin samples from Kratofil et al. (2020) and this study were conducted in a similar manner: skin samples were first dried and homogenized into a fine powder, lipids were extracted because they are $^{13}C$ depleted (DeNiro & Epstein 1977), and samples subsequently processed using a ThermoFinnigan Delta V isotope ratio mass spectrometer (IRMS) coupled with a NA 1500 Ne elemental analyzer. Lipid extractions were done chemically for both Kratofil et al. (2020) samples (with two cell volumes of dichloromethane at 25°C, 500 psi) and the samples analysed herein (following Caputo et al. 2025, with a 2:1 chloroform: methanol mixture). Specific laboratory lipid extraction methods can be found in Kratofil et al. (2020) and Caputo et al. (2025).

Supporting Information S3: Supplemental spatial metric methods & results

*Continuous-time movement model fitting*

Filtered satellite tag location data were fit to continuous-time movement models using the *ctmmUtils* and *ctmm* R packages (Fleming & Calabrese 2023; Johnson & London 2025). Argos error ellipse measurements were incorporated into the model. For tags deployed in 2007, where only location quality class was available, we used corresponding class-specific estimates of ellipsoidal uncertainty from Vincent et al. (2002). Additionally, some tags (n = 3) transmitted both Argos and Fastloc-GPS locations; for these tags, we assigned GPS locations error ellipses based on the number of satellites that were used to estimate the position, following values from Dujon et al. (2014). Any locations on land were rerouted around the 20-meter isobath using the *pathroutr* package (London 2020).

*Resource selection functions*

Stage 1 resource selection functions (RSFs; i.e., individual-level) were fit with environmental variables reflecting seafloor topography and dynamic ocean activity. Correlations between environmental variables were assessed prior to modeling; where two variables were correlated (Pearsons' correlation > 0.5), only one was included for subsequent modeling. In these cases, the variable with the more direct ecological interpretation (e.g., known proxy of upwelling processes) was selected. Details on the variables, their resolution, source, and whether or not they were included in final RSFs are provided in Table S1. Variable selection was not undertaken at the stage 1 RSF level as the goal of the two-stage approach in our study was to assess among-individual variation (and among-cluster variation) in habitat selection of common habitat features.

**Table S1.** Environmental variables and source information for variables considered in resource selection functions (RSFs).

| Variable | Spatial resolution | Temporal resolution | Source | Included/excluded in RSF models |
|---|---|---|---|---|
| Seafloor depth (depth) | 1 km | NA | GEBCO[1] | Included |
| Seafloor slope (slope) | 1 km | NA | GEBCO[1] | Included |

| | | | | |
|---|---|---|---|---|
| Sea surface temperature (sst) | 9 km | Daily | Copernicus Global Ocean Physics Re-analysis[2] | Included |
| Standard deviation of SST (sst_sd) | 27 km | Daily | Calculated across 3 pixels using SST data | Included |
| Sea surface salinity (sss) | 9 km | Daily | Copernicus Global Ocean Physics Reanalysis[2] | Excluded; correlated with ssh |
| Mixed layer depth (mld) | 9 km | Daily | Copernicus Global Ocean Physics Re-analysis[2] | Included |
| Sea surface height (ssh) | 9 km | Daily | Copernicus Global Ocean Physics Re-analysis[2] | Included |
| Standard deviation of SSH (ssh_sd) | 27 km | Daily | Calculated across 3 pixels using SSH data | Included |
| Current magnitude (cmag; derived from u- and v- components of horizontal current velocity) | 9 km | Daily | Copernicus Global Ocean Physics Re-analysis[2] | Excluded; correlated with ssh_sd |
| 30-day lagged Surface chlorophyll-a concentration (logged; chla30) | 4 km | Daily | Copernicus Ocean Color[3]; value extracted for date 30 days prior to date of locations | Included |

[1]https://www.gebco.net/data-products/gridded-bathymetry-data

[2]https://data.marine.copernicus.eu/product/GLOBAL_MULTIYEAR_PHY_001_030/description

[3]https://data.marine.copernicus.eu/product/OCEANCOLOUR_GLO_BGC_L4_MY_009_104/description

*Spatial metric model specifications*

Models examining the influence of social factors on space use metrics are detailed in Table S2. All models were fit in a Bayesian framework using the *brms* package (Bürkner 2018) with 6,000 iterations and a 3,000 warm-up period. We specified diffuse priors for all models as we did not have strong prior knowledge on the parameter distributions (Table S2). For RSF models, habitat variables were centered/scaled prior to model fitting.

For models assessing long-term predictions (i.e., predictions iii in Figure 1, S1), we only included one individual of same-cluster tagged dyads that remained spatially associated during their period of overlap based on findings from the dyadic distances model (i.e., to mitigate pseudoreplication).

The stage 2 RSFs did not include covariates for tag deployment year or duration, as was done for the spatial and core range overlap models. The random effect for tag ID in the stage 2 RSF accounts for any differences due to that individual (including tag deployment year), and the covariance matrices incorporated from stage 1 RSFs will generally reflect differences in among individuals in the amount of data (i.e., deployment length).

Only a subset of tagged dyads (n = 273 dyads) had pairwise genetic relatedness information available and thus separate spatial/core range overlap models were ran for this sub-analysis. Due to limited sample size across unique cluster pairs in the genetic similarity models, a binary predictor for same/different cluster was included in place of unique cluster pair.

**Table S2.** Spatial metric model parameters and priors. Note: stage 2 RSFs (i.e., meta-mixed effects models) were not fit in a Bayesian framework and thus no priors are listed. Parameter types: intercepts (a), slopes (b), distributional parameters (shape, phi, zi).

| Model (family, link function) | Parameter | Prior |
|---|---|---|
| Median distance between dyad (gamma, log link) | a | student_t(3, 4.1, 2.5) |
| | b(HWI) | flat |
| | a(mmid1-id2) | student_t(3, 0, 2.5) |
| | sd(mmid1-id2) | student_t(3, 0, 2.5) |
| | shape | gamma(0.01, 0.01) |
| Spatial range overlap (beta, identity link) | a | student_t(3, 0, 2.5) |
| | phi | gamma(0.01, 0.01) |
| | b(coeffs): cluster pair, same/different year, duration difference (genetic subset: same/different cluster, year, duration difference, genetic relatedness) | flat |
| | a(mmtagid1-tagid2) | student_t(3, 0, 2.5) |
| | sd(mmtagid1-tagid2) | student_t(3, 0, 2.5) |
| Core range overlap (zero-inflated beta, identity link) | a | student_t(3, 0, 2.5) |
| | phi | gamma(0.01, 0.01) |
| | zi | beta(1, 1) |
| | b(coeffs): cluster pair, same/different year, duration difference (genetic subset: same/different cluster, year, duration difference, genetic relatedness) | flat |
| | a(mmtagid1-tagid2) | student_t(3, 0, 2.5) |
| | sd(mmtagid1-tagid2) | student_t(3, 0, 2.5) |
| Stage 1 RSFs (Bernoulli, logit link) | a | student_t(3, 0, 2.5) |
| | b(coeffs): RSF coefficients (Table S1) | flat |
| Stage 2 RSFs (multivariate normal, meta-mixed effects) | a | NA |
| | b(coeffs): cluster | NA |
| | a(tagid) | NA |

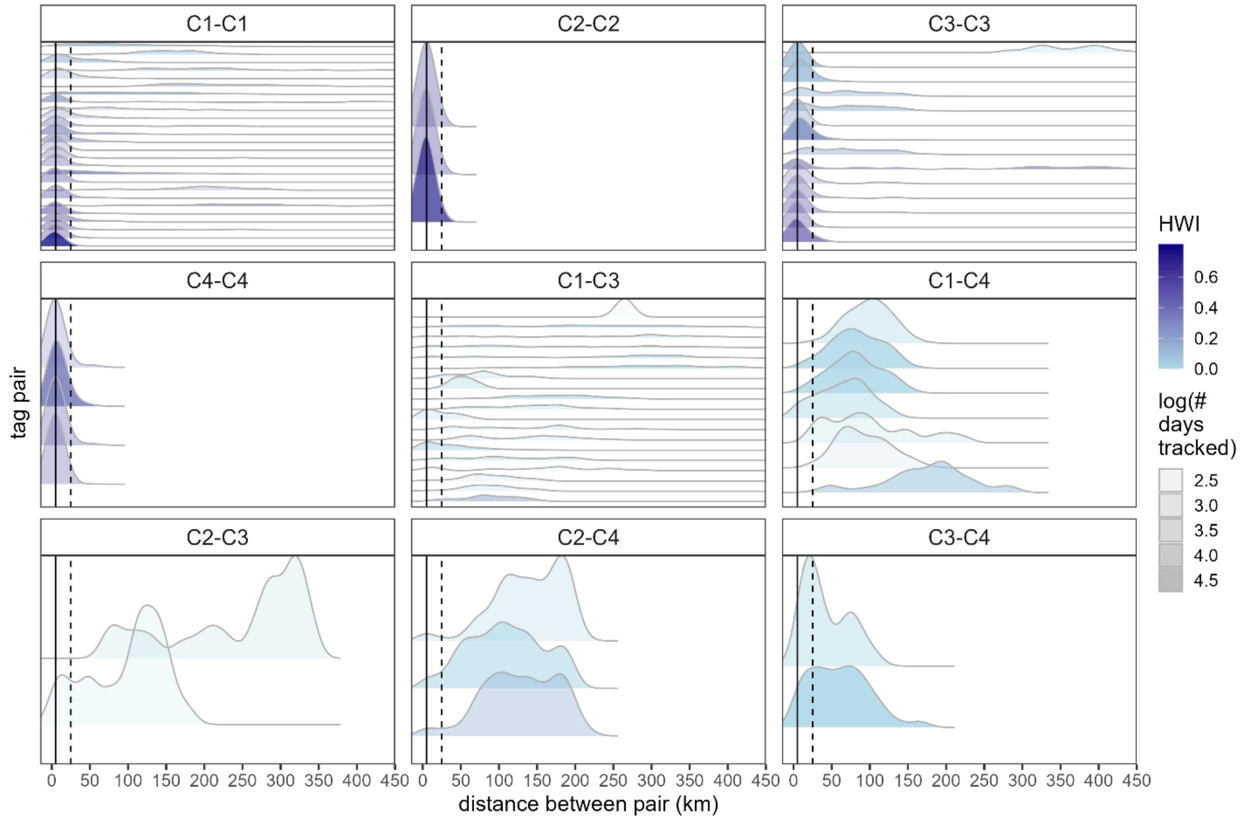


**Figure S2.** Distribution of distances between tagged dyads sorted by decreasing pairwise association strength (half-weight index, HWI) and faceted by cluster pair (i.e., C#-C#). The transparency of density distributions is scaled by the logged number of days tracked for the pair, so that longer tracking durations are less transparent. The solid vertical line represents 5 km, and the dashed vertical line represents 25 km.

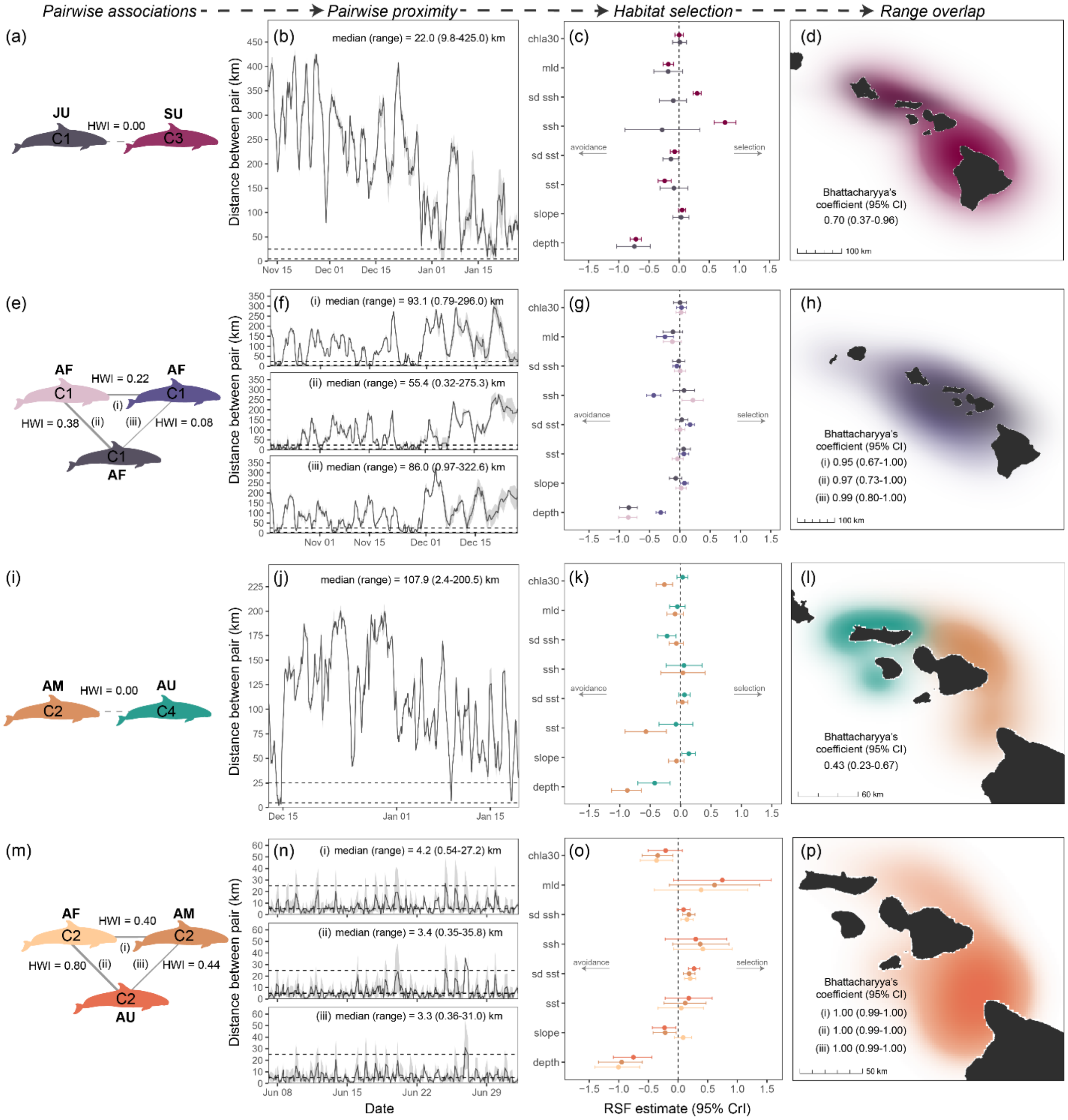


**Figure S3.** Additional cases exhibiting short-term intra- and inter-cluster social associations and space use. Emergence of intra-cluster space use from social bonds (e-h; m-p) from fine to coarse scales (left to right) and between-cluster partitioning (a-c; i-l) in the short-term. Each panel shows (1st column) pairwise association strength (half-weight index, HWI) among the focal individuals (i, ii, iii) with cluster membership (colour and "C#") and age-sex class (A = adult, S

= sub-adult, J = juvenile; F = female, M = male, U = unknown); (2nd column) pairwise proximity (distance between each pair) for the period of overlap, with 5-km and 25-km indicated by horizontal dashed lines; (3rd column) habitat selection coefficients of individuals (estimates and 95% credible intervals, CrI; see Table S1 for variable codes); and (4th column) spatial range overlap, with the Bhattacharyya's coefficient values and 95% confidence intervals (CI) shown for each pair.

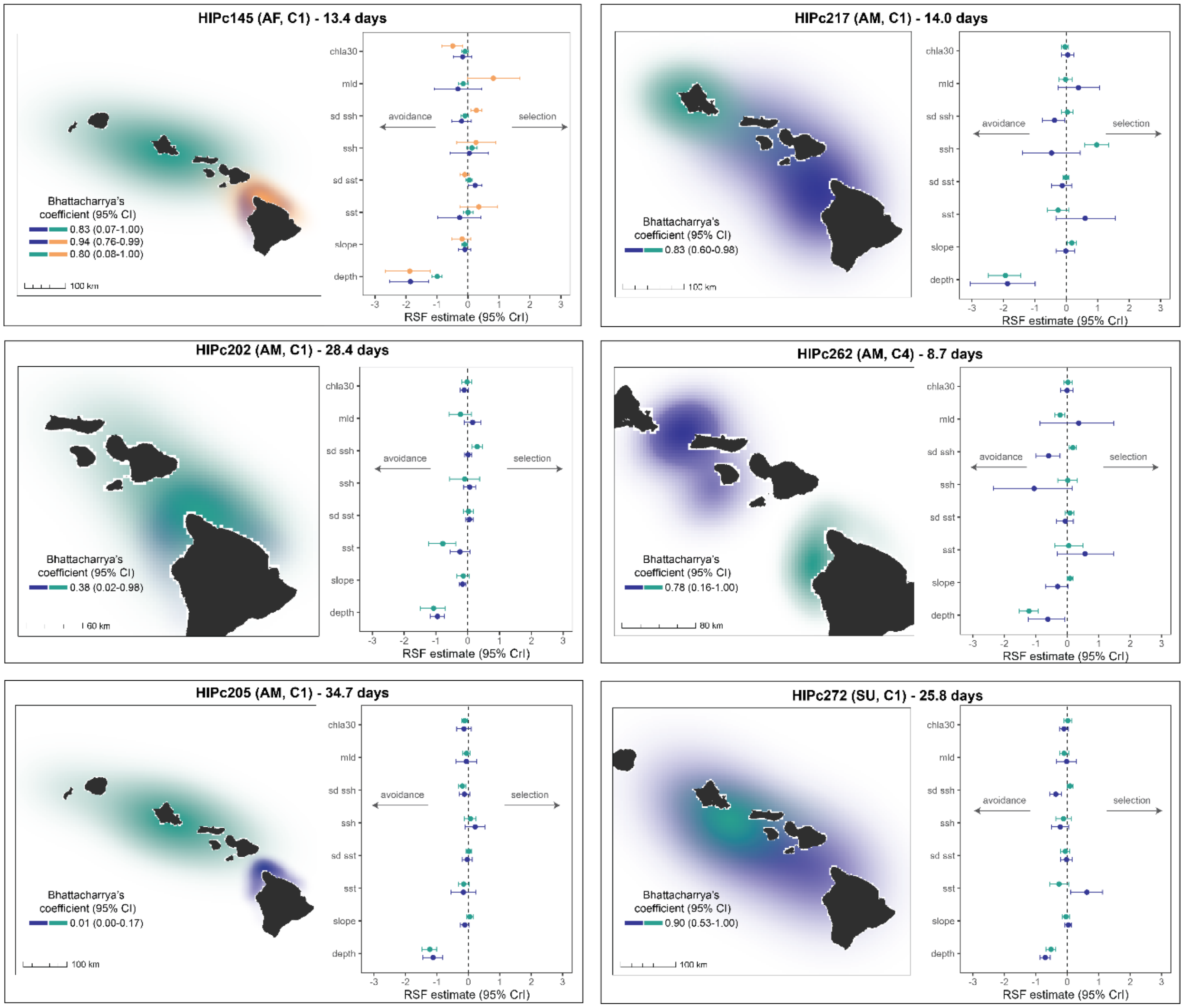


**Figure S4.** Comparative spatial behaviour (spatial range: left sub-panels, resource selection: right sub-panels) of individuals tagged more than once during the study period. Catalogue ID, age/sex class (A = adult, S = sub-adult, F = female, M = male), cluster membership (C#) and duration are indicated within each panel. Overlap between spatial ranges (Bhattacharyya's coefficient) and associated confidence intervals (CI) are provided in each map. Resource selection function coefficient names are provided in Table S1.

**Table S3.** Spatial metric model parameter estimates. Coefficient estimates (i.e., b parameters) that have 95% credible intervals (CrI) excluding zero are shown in bold. See Table S2 for parameter definitions. For models including coefficients for unique cluster pair (i.e., CX-CY), the reference level was set to C1-C1; C1 was set as the reference level for models including cluster. For the stage 2 RSF model, confidence intervals (CR) are provided.

| Model (sample size) | Parameter | Estimate | Lower 95% CrI | Upper 95% CrI |
|---|---|---|---|---|
| Median distance between dyads (n=80 dyads) | a | 4.56 | 4.06 | 5.06 |
| | **b(HWI)** | **-4.51** | **-5.64** | **-3.36** |
| | sd(mmid1-id2) | 1.26 | 0.89 | 1.70 |
| | shape | 2.09 | 1.35 | 3.04 |
| Spatial range overlap (n=1,423 dyads) | a | 0.96 | 0.51 | 1.40 |
| | b(C1-C2) | -0.09 | -0.69 | 0.52 |
| | b(C1-C3) | 0.14 | -0.24 | 0.53 |
| | **b(C1-C4)** | **-1.30** | **-1.88** | **-0.72** |
| | b(C2-C2) | 0.59 | -0.71 | 1.94 |
| | b(C2-C3) | 0.00 | -0.77 | 0.78 |
| | **b(C2-C4)** | **-1.46** | **-2.40** | **-0.50** |
| | b(C3-C3) | 0.31 | -0.46 | 1.07 |
| | b(C3-C4) | -0.70 | -1.45 | 0.05 |
| | b(C4-C4) | -1.17 | -2.45 | 0.13 |
| | **b(same_year = TRUE)** | **0.53** | **0.31** | **0.75** |
| | **b(duration_diff)** | **-0.10** | **-0.18** | **-0.02** |
| | sd(mmtagid1-tagid2) | 1.19 | 0.97 | 1.47 |
| | phi | 3.86 | 3.58 | 4.14 |
| Core range overlap (n=1,423 dyads) | a | -0.60 | -0.85 | -0.36 |
| | b(C1-C2) | -0.20 | -0.55 | 0.16 |
| | b(C1-C3) | -0.13 | -0.37 | 0.11 |
| | **b(C1-C4)** | **-0.55** | **-0.92** | **-0.18** |
| | b(C2-C2) | 0.32 | -0.49 | 1.14 |
| | b(C2-C3) | -0.39 | -0.85 | 0.07 |
| | **b(C2-C4)** | **-0.86** | **-1.67** | **-0.10** |
| | b(C3-C4) | -0.29 | -0.74 | 0.13 |
| | **b(C4-C4)** | **-0.54** | **-1.01** | **-0.07** |
| | **b(same_year = TRUE)** | **1.02** | **0.05** | **1.99** |
| | **b(duration_diff)** | **0.26** | **0.05** | **0.47** |
| | sd(mmtagid1-tagid2) | 0.59 | 0.45 | 0.77 |
| | phi | 3.86 | 3.55 | 4.19 |
| | zi | 0.26 | 0.24 | 0.29 |
| Spatial range overlap – genetic relatedness subset (n=273 dyads) | a | 0.78 | 0.18 | 1.38 |
| | b(same_clust = TRUE) | 0.22 | -0.02 | 0.47 |
| | **b(same_year = TRUE)** | **0.55** | **0.05** | **1.07** |
| | b(duration_diff) | -0.03 | -0.29 | 0.23 |
| | b(genetic_relatedness) | -0.20 | -1.43 | 1.08 |

| | | | | |
|---|---|---|---|---|
| | sd(mmtagid1-tagid2) | 1.40 | 1.01 | 1.94 |
| | phi | 4.53 | 3.80 | 5.33 |
| Core range overlap – genetic relatedness subset (n=273 dyads) | a | -0.96 | -1.38 | -0.55 |
| | b(same_clust = TRUE) | 0.21 | -0.07 | 0.49 |
| | b(same_year = TRUE) | 0.22 | -0.32 | 0.74 |
| | b(duration_diff) | -0.10 | -0.33 | 0.12 |
| | b(genetic_relatedness) | -0.18 | -1.64 | 1.24 |
| | sd(mmtagid1-tagid2) | 0.87 | 0.54 | 1.30 |
| | phi | 4.70 | 3.81 | 5.68 |
| | zi | 0.24 | 0.19 | 0.30 |
| | ***Stage 2 RSF meta-mixed effects regression*** | | | |
| **RSF variable** | **Parameter** | **Estimate** | **Lower 95% CI** | **Upper 95% CI** |
| depth | a | -0.94 | -1.04 | -0.84 |
| | b(C2) | 0.07 | -0.20 | 0.34 |
| | **b(C3)** | **0.33** | **0.17** | **0.50** |
| | b(C4) | 0.23 | -0.03 | 0.50 |
| slope | a | -0.02 | -0.05 | 0.003 |
| | b(C2) | -0.04 | -0.11 | 0.03 |
| | b(C3) | -0.01 | -0.05 | 0.04 |
| | b(C4) | 0.03 | -0.04 | 0.10 |
| sst | a | 0.03 | -0.06 | 0.13 |
| | b(C2) | -0.10 | -0.33 | 0.13 |
| | b(C3) | -0.04 | -0.20 | 0.11 |
| | b(C4) | -0.05 | -0.29 | 0.19 |
| sst_sd | a | 0.05 | 0.02 | 0.09 |
| | b(C2) | 0.001 | -0.09 | 0.09 |
| | b(C3) | -0.03 | -0.09 | 0.03 |
| | b(C4) | -0.03 | -0.12 | 0.06 |
| ssh | a | -0.02 | -0.14 | 0.09 |
| | b(C2) | 0.09 | -0.19 | 0.38 |
| | **b(C3)** | **0.20** | **0.01** | **0.38** |
| | b(C4) | 0.01 | -0.27 | 0.30 |
| ssh_sd | a | -0.04 | -0.10 | 0.02 |
| | b(C2) | 0.15 | -0.004 | 0.30 |
| | b(C3) | 0.07 | -0.03 | 0.17 |
| | b(C4) | 0.02 | -0.14 | 0.18 |
| mld | a | -0.002 | -0.07 | 0.07 |
| | b(C2) | 0.12 | -0.04 | 0.28 |
| | b(C3) | -0.04 | -0.15 | 0.06 |
| | b(C4) | -0.04 | -0.20 | 0.11 |
| chla30 | a | -0.09 | -0.14 | -0.04 |
| | b(C2) | -0.12 | -0.24 | 0.004 |
| | b(C3) | 0.03 | -0.05 | 0.10 |
| | b(C4) | 0.08 | -0.04 | 0.19 |

Supporting Information S4: Supplemental stable isotopes data analysis methods & results

*Isotopic summaries, temporal and demographic effects*

Prior to formal data analyses, carbon isotope values were Suess-corrected using the *SuessR* R package (Clark *et al.* 2021) to account for anthropogenic-driven changes in baseline ocean carbon levels over the 12-year sampling period. This package includes built-in regional baseline values to inform correction factors; we used the Aleutian Islands region to Suess-correct our samples as the carbon levels are the same for the Hawaiian Islands (Velasquez-Vacca *et al.* 2024). We corrected values to the year of the most recent samples (2020) so that values reported herein will be comparable to other contemporary studies in Hawai‘i.

We assessed potential temporal and demographic effects on $\delta^{13}$C and $\delta^{15}$N values using Bayesian generalised linear regression models in the *brms* package (Bürkner 2018). Out of the 80 total samples, 14 represented individuals sampled twice (i.e., seven individuals) over the study period; for analytical purposes, we considered these samples to be independent given they were obtained in separate years. Separate models were fit for $\delta^{13}$C and $\delta^{15}$N values assuming a Gaussian error distribution and identity link function, and considering sample year, sample season, age class, sex, and social cluster as fixed effects. We aggregated sampling year into two sampling periods to attempt to alleviate uneven sample sizes across years (2008-2011, n = 54 samples; 2015-2020, n = 26 samples). Samples were categorised into oceanographic seasons (Flament 1996) based on one month prior to that of sampling to reflect the season during which foraging is reflected by the sample (i.e., approximately one month tissue turnover rate; Giménez *et al.* 2016). Due to low sample size during the winter compared to all other seasons, we combined samples into summer/fall (n = 43 samples) and winter/spring (n = 37 samples) seasons. Similarly, there were limited juveniles (n = 12) and sub-adults sampled (n = 9) compared to adults (n = 59), and thus juveniles and sub-adults were combined into one category. Sample size by sex was less skewed (males, n = 33; females, n = 47). Models were fit with four chains with 6,000 iterations and a 3,000-iteration warm-up phase. Model convergence was ensured through $\hat{R}$ values (<1.05), mixing of chains, and posterior predictive checks.

There was evidence for an effect of sampling period, with samples from 2015-2020 having higher $\delta^{13}$C and $\delta^{15}$N values (posterior estimates = 0.48, 0.50, 95% CrIs = 0.29 to 0.67, 0.22 to 0.79, respectively) compared to 2008-2011 samples; however, sample size between periods was skewed (n = 54, 26; Table S4) and the difference in mean values was within 1‰ (Figure S5). Samples from summer/fall season had lower $\delta^{15}$N values compared to winter/spring (posterior estimate = -0.61, 95% CrI = -0.97 to -0.24), although the observed difference was less than 1‰; there was no evidence for an effect of season on $\delta^{13}$C values (Table S4). Sub-adults/juveniles had higher $\delta^{13}$C values than adults (posterior estimate = 0.23, 95% CrI = 0.03 to 0.44), although sample size was skewed (Table S4; Figure S5). There was no estimated effect of age class on $\delta^{15}$N values, or sex on $\delta^{13}$C or $\delta^{15}$N values (Table S4).

**Table S4.** Bayesian generalised linear model parameter estimates for $\delta^{13}$C and $\delta^{15}$N in relation to temporal and demographic covariates. Covariates with a 95% credible interval (CrI) that crosses zero are bolded.

| Model | Parameter | Estimate | Lower 95% CrI | Upper 95% CrI |
|---|---|---|---|---|
| $\delta^{13}$C | a | -16.35 | -16.52 | -16.19 |
| | **b(Period: 2015-2020)** | **0.48** | **0.29** | **0.67** |
| | b(Season: summer-fall) | 0.20 | -0.04 | 0.45 |
| | b(Cluster 2) | -0.15 | -0.36 | 0.06 |

| | | | | |
|---|---|---|---|---|
| | b(Cluster 3) | -0.04 | -0.34 | 0.26 |
| | b(Cluster 4) | 0.00 | -0.26 | 0.26 |
| | **b(Age class: sub-adult/juv)** | **0.23** | **0.04** | **0.43** |
| | b(Sex: male) | -0.01 | -0.19 | 0.16 |
| $\delta^{15}N$ | a | 12.33 | 12.09 | 12.58 |
| | **b(Period: 2015-2020)** | **0.50** | **0.22** | **0.78** |
| | **b(Season: summer-fall)** | **-0.61** | **-0.97** | **-0.25** |
| | b(Cluster 2) | 0.00 | -0.31 | 0.31 |
| | b(Cluster 3) | 0.25 | -0.19 | 0.68 |
| | b(Cluster 4) | -0.14 | -0.52 | 0.25 |
| | b(Age class: sub-adult/juv) | 0.03 | -0.27 | 0.32 |
| | b(Sex: male) | 0.06 | -0.19 | 0.32 |

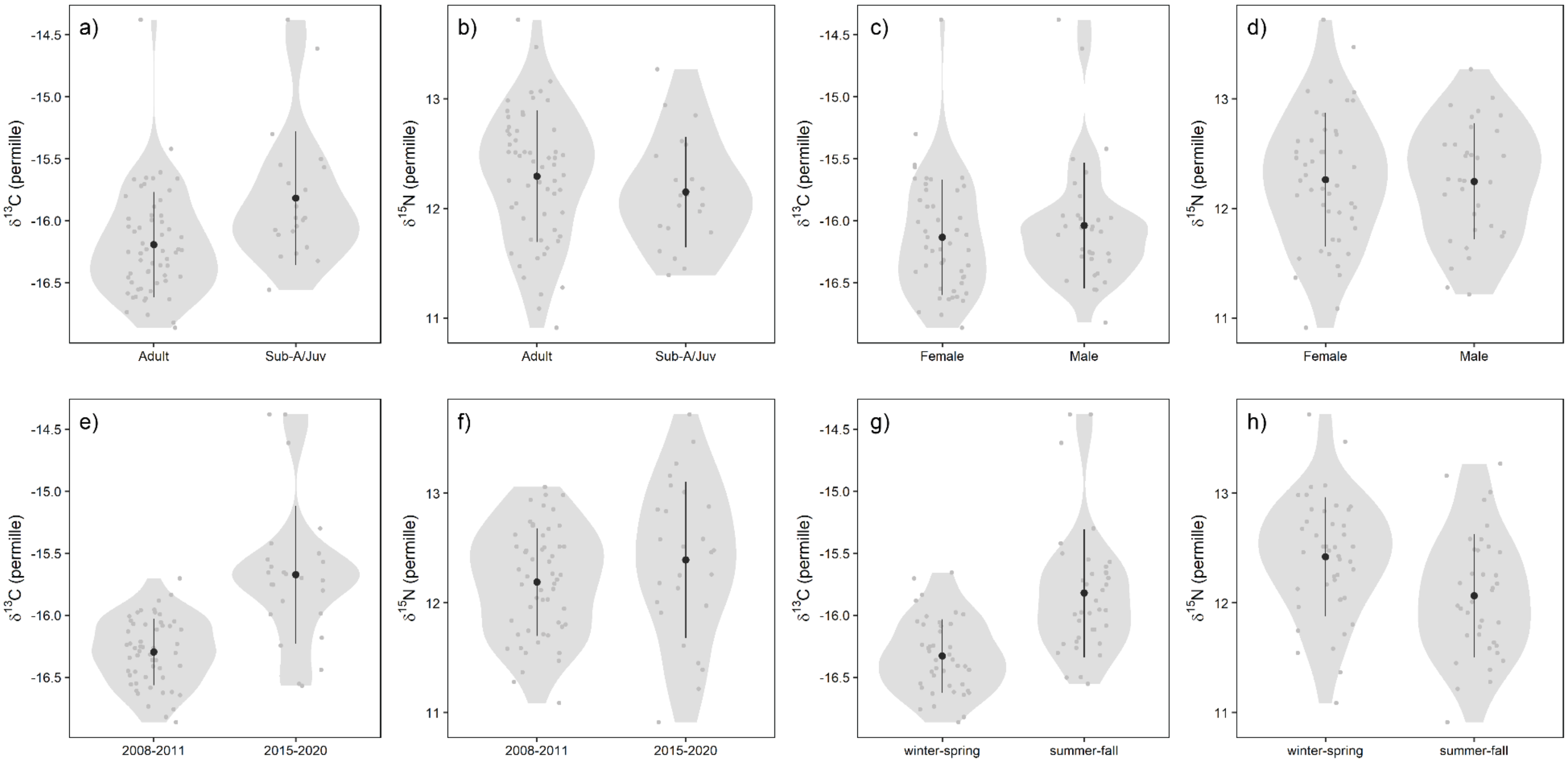


**Figure S5.** Violin plots of $\delta^{13}$C and $\delta^{15}$N values by (a, b) age class, (c, d) sex, (e, f) sampling period, and (g, h) sampling season, respectively. Points represent the mean value, and lines represent the standard deviation of the mean. Observations are shown as jittered points.

*Cluster-level isotopic niches*

Cluster-level isotopic niches were quantified in a Bayesian framework using the *nicheROVER* R package (Lysy *et al.* 2023; Swanson *et al.* 2015). For each social cluster, we fit a single model with 10,000 Markov chain Monte Carlo (MCMC) iterations and a flat Normal-Inverse-Wishart prior (Lysy *et al.* 2023). Niche size was considered the region of the posterior distribution with a 95% probability of containing individuals from each social cluster (Swanson *et al.* 2015). Pairwise niche overlap was assessed as the percentage probability of an individual from one social cluster falling within the niche space of another social cluster's niche (Swanson *et al.* 2015). We quantified the posterior distribution of niche overlap for both the overall niche space (95% probability region) and the core niche space (40% probability region) among social clusters (Table S5; Swanson *et al.* 2015).

**Table S5.** Posterior probability of bivariate $\delta^{13}C$ and $\delta^{15}N$ isotopic niche overlap (with 95% credible intervals) among social clusters for the overall niche (95% region) and core niche (40% region).

| | | **Mean probability of overlap (95% credible interval)** | | | |
|---|---|---|---|---|---|
| | | Cluster 1 | Cluster 2 | Cluster 3 | Cluster 4 |
| **Overall (95% region)** | Cluster 1 | - | 86.0 (68.5-97.5) | 87.8 (65.6-99.0) | 51.9 (32.6-72.5) |
| | Cluster 2 | 83.1 (65.0-96.3) | - | 71.8 (41.7-94.9) | 41.3 (21.8-64.3) |
| | Cluster 3 | 70.7 (45.8-92.9) | 54.3 (27.7-84.6) | - | 47.8 (28.3-71.4) |
| | Cluster 4 | 92.3 (76.1-99.7) | 90.9 (68.6-99.8) | 95.5 (80.0-100.0) | - |
| **Core (40% region)** | Cluster 1 | | 33.3 (19.1-51.0) | 29.8 (11.5-53.9) | 13.1 (6.4-22.6) |
| | Cluster 2 | 30.7 (17.7-47.2) | | 14.5 (2.5-35.1) | 9.1 (3.5-17.4) |
| | Cluster 3 | 20.9 (9.6-38.3) | 12.6 (3.4-27.6) | | 12.6 (6.0-22.6) |
| | Cluster 4 | 36.4 (13.0-63.4) | 21.5 (3.1-53.0) | 54.3 (27.5-80.7) | |

*Mixing models: prey data, methods, and sensitivity analyses*

We completed a literature search and compiled bulk $\delta^{13}$C and $\delta^{15}$N values available for 13 of 19 known false killer whale prey species that have been identified from observations of feeding events or stomach contents (Table S6; K. West Unpublished; Baird *et al.* 2008, 2021; Zaeschmar & Baird 2025). The six species that were unaccounted for either had another species in their functional group with stable isotope data (e.g., cephalopods, pelagic-oceanic fishes, neritic reef fish) or current evidence suggests they comprise a very small proportion of false killer whales' diet (elasmobranchs; Zaeschmar & Baird 2025). Thus, our available data were reasonably representative of their known diet.

Prey $\delta^{13}$C isotope values were Suess-corrected following the same methods described above for the false killer whale samples, particularly to account for temporal changes in anthropogenic $CO^2$ emissions. Studies accounted for the potential effect of lipids on $\delta^{13}$C either through chemical extraction (Carlisle *et al.* 2021; Graham 2007) or mathematical correction (Blum *et al.* 2013; Choy *et al.* 2015), while two studies reported low lipid content removing the necessity of lipid extraction (squid, Gloeckler *et al.* 2018; reef-associated fish, Sackett *et al.* 2015). There were several prey species with multiple studies reporting stable isotope values. For some of these studies, raw stable isotope values were made available, while others only reported means and standard deviations (SDs) for each prey species. Therefore, to comprehensively integrate values from all relevant studies, we resampled 1,000 values from the distributions of $\delta^{13}$C and $\delta^{15}$N for each prey/study and combined them into a cumulative distribution (and single mean and SD) for each prey species following Carlisle et al. (2021).

To discern whether differences in stable isotope methods across studies could potentially confound mixing model results, we ran a second version of the mixing model using only single-study prey data (i.e., data for each species comes from only one study). These studies also sampled prey during the same general time period (except for reef-associated fish), which mitigates any large-scale interannual variation in isotope values. We used values from Gloeckler et al. (2018) for mesopelagic cephalopods (purpleback flying squid, *Sthenoteuthis oualaniensis*, sampled in 2011 and 2014), Sackett et al. (2015) for neritic reef fish (uku, kahala, ulua; sampled 2003-2004 (uku, kahala) and 2009 (ulua)), and Carlisle et al. (2021) for all other prey species listed in Table S6 (sampled in 2011-2013).

**Table S6.** Summary of known false killer whale prey species sampled in Hawaiian waters or Central North Pacific region more broadly. Sample size (n*) reflects the number of samples from the referenced studies combined. Mean and standard deviation (SD) isotope values reflect the cumulative distribution of raw and resampled isotope distributions. Carbon isotope values ($\delta^{13}C$) were Suess-corrected to the year 2020 prior to summarizing the data. DVM = diel vertical migration.

| Hawaiian name (common name; scientific name) | n* all / n single | Mean (SD) $\delta^{13}C$ | Mean (SD) $\delta^{15}N$ | Evidence for DVM | Mean (SD) trophic position | Habitat zone: vertical/horizontal | Data, trophic level, and habitat references § |
|---|---|---|---|---|---|---|---|
| A‘u ku (broadbill swordfish; *Xiphias gladius*) | 28/9 | -17.6 (1.7) | 13.1 (1.7) | X | 4.9 (0.9) | Epi-mesopelagic/oceanic | 1,2,3 / 3,7,8 |
| A‘u (shortbill spearfish; *Tetrapturus angustirostris*) | 8/8 | -17.2 (0.7) | 11.3 (2.1) | | 4.5 (0.8) | Epipelagic/oceanic | 2 / 9,10,22 |
| ‘Ahi po‘onui (bigeye tuna; *Thunnus obesus*) | 55/9 | -16.9 (0.7) | 11.3 (1.1) | X | 5.0 (0.6) | Epi-mesopelagic/oceanic | 1,2,3,5 / 3,11 |
| Ono (wahoo; *Acanthocybium solandri*) | 11/11 | -17.4 (0.4) | 12.0 (1.4) | | 4.3 (0.2) | Epipelagic/oceanic | 2 / 12,22 |
| Mahimahi (dolphinfish; *Coryphaena hippurus*) | 30/9 | -16.7 (1.1) | 10.7 (1.6) | | 4.3 (0.5) | Epipelagic/oceanic | 1,2,3 / 3,13 |
| ‘Ahi (yellowfin tuna; *Thunnus albacares*) | 30/9 | -16.5 (1.0) | 9.8 (1.9) | X | 4.6 (0.1) | Epipelagic/oceanic | 1,2,3,5 / 3,14,15 |
| Aku (skipjack tuna; *Katsowonus pelamis*) | 24/12 | -17.0 (0.8) | 9.9 (1.8) | X | 4.3 (0.5) | Epipelagic/oceanic | 1,2,3 / 3,16 |
| ‘Ahi palaha (albacore tuna; *Thunnus alalunga*) | 3/3 | -18.5 (0.4) | 11.4 (0.6) | X | 4.3 (0.2) | Epi-mesopelagic/oceanic | 2 / 17,22 |
| Opah (moonfish; *Lampris guttatus*) | 40/10 | -18.6 (1.1) | 11.7 (1.0) | X | 4.5 (0.1) | Mesopelagic/oceanic | 1,2,3 / 3,18 |
| Purpleback flying squid (*Sthenoteuthis oualaniensis*) † | 28/3 | -18.6 (0.5) | 9.7 (2.1) | X | 2.9 (NA) | Meso-bathypelagic/oceanic | 2,4,25 / 19 |

| | | | | | | |
|---|---|---|---|---|---|---|
| Uku (blue-green snapper; *Aprion virescens*) | 24/24 | -16.3 (0.5) | 9.5 (0.5) | 4.3 (0.2) | Benthopelagic/coastal | 6 / 20,21,23,24 |
| Kahala (amber, almaco jack; *Seriola dumerili, rivoliana*) | 8/8 | -16.7 (0.2) | 11.2 (0.2) | 4.4 (0.03) | Benthopelagic/coastal | 6 / 23,24 |
| Ulua (giant trevally; *Caranx ignobilis*) | 8/8 | -13.7 (1.3) | 11.5 (1.4) | 3.8 (0.3) | Benthic/coastal | 6 / 23,24 |

* Sample size reflects all available samples (i.e., all data references) and single studies of prey data (“single”) used in the sensitivity analyses

§ References: 1 Blum et al. (2013); 2 Carlisle et al. (2021); 3 Choy et al. (2015); 4 Gloeckler et al. (2018); 5 Graham (2007); 6 Sackett et al. (2015); 7 Abecassis et al. (2012); 8 Dewar et al. (2011); 9 Arostegui et al. (2019); 10 Arostegui et al. (2024); 11 Musyl et al. (2003); 12 Sepulveda et al. (2011); 13 Whitney et al. (2016); 14 Brill et al. (1999); 15 Lam et al. (2020); 16 Shaefer & Fuller (2007); 17 Domokos et al. (2007); 18 Polovina et al. (2008); 19 Jereb & Roper (2010); 20 Tanaka et al. (2022); 21 Asher et al. (2017); 22 FishBase (2025); 23 Sackett et al. (2017); 24 HI DLNR; 25 Carlisle et al. (2015)

† Sample size for purpleback flying squid: one of the referenced studies (26) notes a sample size of 25 for squid with carbon and 127 for nitrogen, and we conservatively report the sample size of 25 (n = 28 with data from an additional reference) here.

We aggregated the 13 prey sources into groups *a posteriori* to increase the discrimination power and interpretability of the mixing model (Phillips *et al.* 2005, 2014; Stock *et al.* 2018). We defined four prey groups based largely on ecological and functional traits (Table S7): epipelagic predatory fishes (shortbill spearfish/a‘u, wahoo/ono, mahimahi, yellowfin tuna/‘ahi, skipjack tuna/aku, almaco jack/kahala, blue-green snapper/uku); mesopelagic predatory fishes (broadbill swordfish/a‘u ku, bigeye tuna/‘ahi po‘onui, moonfish/opah, albacore tuna/‘ahi palaha); reef-associated predatory fish (giant trevally/ulua), and mesopelagic cephalopods, using purpleback flying squid as model species. Mixing models were fit using the *MixSIAR* R package following the specifications detailed in the main text (Stock *et al.* 2018; Stock & Semmens 2016). Trophic discrimination factors (TDFs) are incorporated into mixing models to account for discrimination in isotopes between the consumer and prey. We used the TDF values for carbon and nitrogen derived from a related species (bottlenose dolphins, Tursiops truncatus; $\delta^{13}C = 1.01 \pm 0.37‰$, $\delta^{15}N = 1.57 \pm 0.52‰$; Giménez *et al.* 2016) for our model on false killer whales, the closest species based on phylogeny and with a comparable metabolism (McGowen *et al.* 2020). Adequacy of prey sources and the TDF in capturing the consumers (i.e., false killer whales’) diet was assessed using simulated mixing polygons (Smith *et al.* 2013) and visually inspecting tracer plots (i.e., bivariate isotopic space with TDF corrections applied). Simulated mixing polygons and tracer plots indicated that all false killer whale isotope values fell within the isotopic space defined by the prey species with TDFs incorporated (Figures S6, S7). Posterior proportions of groups (and prey within those groups) for each social cluster and the population are provided in Table S8.

**Table S7.** Summary of false killer whale prey groups summarized *a posteriori* in mixing models.

| Group | Species | Description |
|---|---|---|
| Epipelagic predatory fish | a‘u, ono, mahimahi, ‘ahi, aku, kahala, uku | Primarily occur and forage in the epipelagic zone; associate with surface mixed layer; limited diel vertical migrations; oceanic; includes benthopelagic fish |
| Mesopelagic predatory fish | a‘u ku, ‘ahi po‘onui, opah, ‘ahi palaha, | Primarily occur and forage in the mesopelagic zone, but some may migrate to the epipelagic zone over the diel cycle; oceanic |
| Reef-associated predatory fish | ulua | Primarily occur and forage in benthic or benthopelagic zones in shallow coastal habitats |
| Mesopelagic cephalopods | purpleback flying squid | Cephalopods that primarily occur and forage in the meso-bathypelagic zone, known to undertake diel vertical migrations |

Two species—kahala and uku—overlap in habitat with ulua (separate group representing reef-associated game fish) but also use pelagic waters to an extent and isotopically were confounded with several epipelagic species (Figure S7, Table S8). Therefore, these species were grouped with epipelagic predatory fish in the model presented in the main text while ulua was kept separate (most isotopically distinct). To examine the effect of this decision, we also assessed outputs from a simplified mixing space with three prey sources: (1) epipelagic predatory fishes (as in Table S7), (2) shallow and deep reef-associated predatory fish (kahala, uku, and ulua), and (3) mesopelagic prey (mesopelagic predatory fishes from Table S7 combined with mesopelagic cephalopods). We acknowledge that aggregating 13 species into only three to four groups results in coarser depictions of sources to false killer whales' diet, and that some groups include prey species with slightly variable ecology or trophic level. However, we find these groupings still informative for our research questions as they are largely based on vertical and horizontal foraging habitat use of prey.

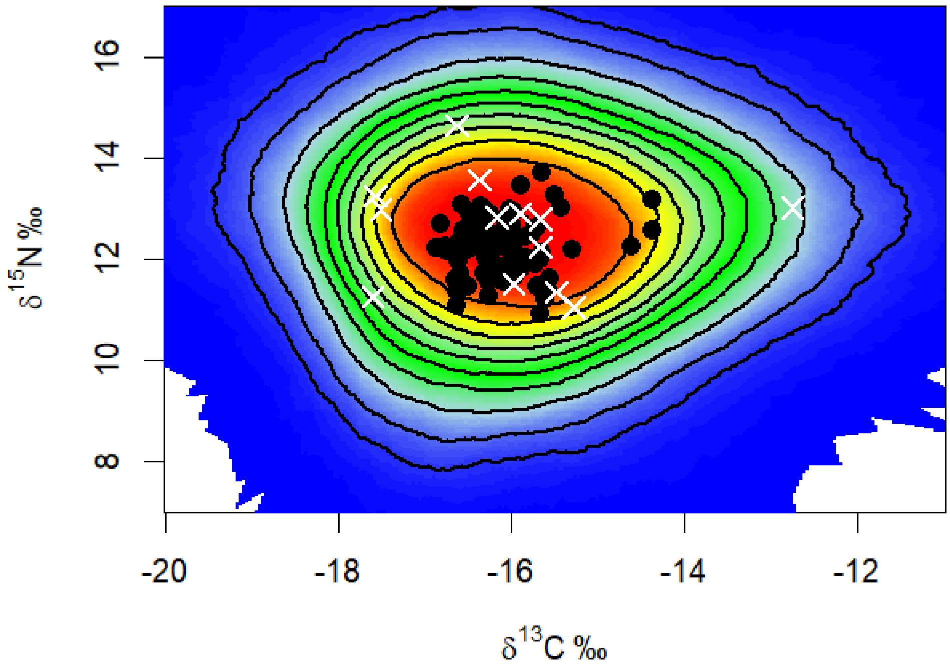


**Figure S6.** Simulated mixing polygon of prey species (white x's) and false killer whales (black dots). The probability that the prey fall within the consumer diet (i.e., false killer whales') increases with each inwards contour line (10% lines).

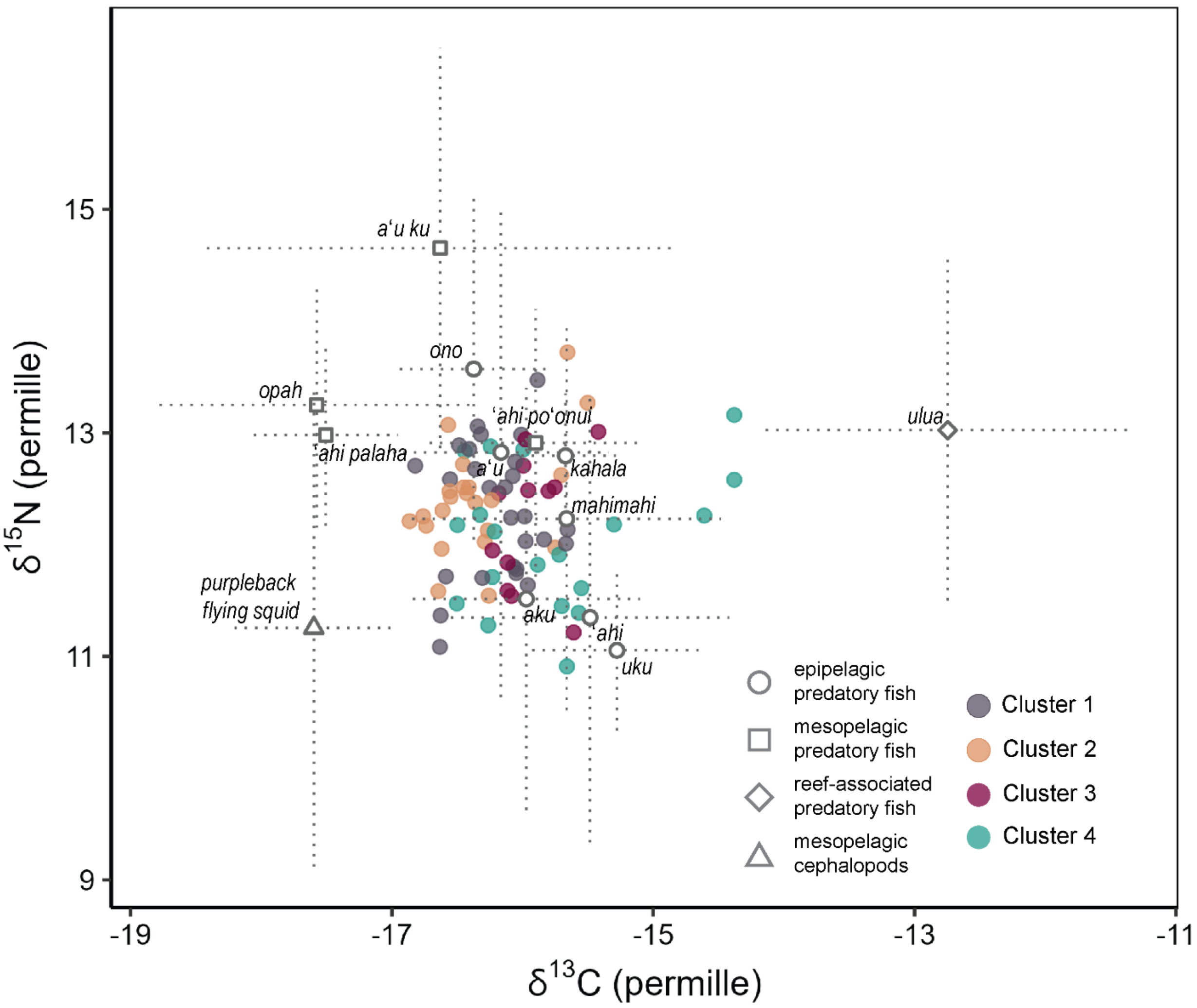


**Figure S7.** Bivariate $\delta^{13}$C and $\delta^{15}$N isotopic space of sampled false killer whales (points) and their prey (white shapes = means, dotted lines = standard deviations; trophic discrimination factors are incorporated).

**Table S8.** Stable isotope mixing model results with posterior mean (95% credible interval; CrI) predicted proportion of each prey group and individual prey species to false killer whales' diet for the population and by social cluster.

| | Posterior mean proportion (95% CrI) | | | | |
|---|---|---|---|---|---|
| **Prey group,** *species* | **Population** | **Cluster 1** | **Cluster 2** | **Cluster 3** | **Cluster 4** |
| **Epipelagic predatory fish** | 0.611 (0.385-0.792) | 0.600 (0.350-0.817) | 0.538 (0.112-0.755) | 0.658 (0.390-0.880) | 0.715 (0.393-0.921) |
| *Mahimahi* | 0.092 (0.001-0.286) | 0.072 (0.001-0.250) | 0.081 (0.001-0.280) | 0.081 (0.001-0.293) | 0.175 (0.001-0.616) |
| *'Ahi* | 0.105 (0.005-0.298) | 0.089 (0.003-0.263) | 0.081 (0.002-0.258) | 0.106 (0.002-0.320) | 0.167 (0.002-0.551) |
| *Ono* | 0.072 (0.003-0.219) | 0.084 (0.001-0.323) | 0.074 (0.001-0.286) | 0.074 (0.001-0.277) | 0.045 (0.001-0.163) |
| *A'u* | 0.068 (0.003-0.223) | 0.078 (0.001-0.300) | 0.066 (0.001-0.266) | 0.073 (0.002-0.289) | 0.050 (0.001-0.231) |
| *Aku* | 0.087 (0.004-0.267) | 0.092 (0.002-0.318) | 0.078 (0.002-0.263) | 0.094 (0.002-0.319) | 0.086 (0.001-0.446) |
| *Uku* | 0.107 (0.002-0.315) | 0.095 (0.001-0.286) | 0.084 (0.001-0.261) | 0.124 (0.001-0.364) | 0.136 (0.001-0.568) |
| *Kahala* | 0.082 (0.003-0.255) | 0.091 (0.001-0.339) | 0.075 (0.002-0.267) | 0.106 (0.002-0.398) | 0.056 (0.001-0.223) |
| **Mesopelagic predatory fish** | 0.256 (0.108-0.440) | 0.261 (0.067-0.475) | 0.335 (0.107-0.551) | 0.221 (0.060-0.431) | 0.177 (0.040-0.372) |
| *'Ahi palaha* | 0.093 (0.004-0.245) | 0.106 (0.002-0.302) | 0.145 (0.002-0.398) | 0.075 (0.002-0.232) | 0.053 (0.001-0.179) |
| *'Ahi po'onui* | 0.068 (0.002-0.214) | 0.071 (0.001-0.272) | 0.068 (0.001-0.266) | 0.072 (0.001-0.276) | 0.054 (0.001-0.251) |
| *A'u ku* | 0.038 (0.002-0.130) | 0.030 (0.001-0.109) | 0.038 (0.001-0.152) | 0.029 (0.001-0.106) | 0.031 (0.001-0.156) |
| *Opah* | 0.058 (0.002-0.178) | 0.054 (0.001-0.182) | 0.084 (0.001-0.291) | 0.044 (0.001-0.156) | 0.039 (0.001-0.151) |
| **Reef-associated predatory fish** (*Ulua)* | 0.047 (0.001-0.151) | 0.034 (0.001-0.116) | 0.031 (0.001-0.108) | 0.047 (0.001-0.156) | 0.053 (0.000-0.231) |
| **Mesopelagic cephalopods** (*Purpleback flying squid)* | 0.086 (0.004-0.227) | 0.104 (0.003-0.269) | 0.096 (0.002-0.274) | 0.074 (0.002-0.227) | 0.055 (0.001-0.202) |

Mixing model outputs using single-study prey isotope values generally had similar findings as the mixing models that used prey isotope values from all available studies. Estimated proportions of the four prey sources (Table S9) in the single-study prey isotope model were close or equal to the proportions estimated by the model using all prey isotope values (Table S8). The biggest differences were slightly increased proportions of reef-associated predatory fish for all clusters (differences in mean posterior proportions between 0.4-0.9) and higher epipelagic predatory fish proportions for Cluster 4 in the single-study model (difference in mean posterior proportion = 0.13; Table S9). Further, kahala and uku had the highest estimated proportions within the epipelagic group, albeit with higher uncertainty (Table S9). These observed differences are likely driven by sample sizes per prey species in the mixing model. The model fitted with single-study prey data had lower sample sizes of most epipelagic and mesopelagic predatory fishes that were more similar to, or even less than, the sample size of the reef-associated predatory fish (Table S6). The *MixSIAR* package incorporates uncertainty in prey source sample size into the mixing models, and thus prey source isotope summaries with higher sample sizes will have less uncertainty compared to those with lower sample sizes (Stock *et al.* 2018; Stock & Semmens 2016). Additionally, the isotopic signatures of some epipelagic predatory fish that had higher proportions in the all-prey isotope value model (e.g., mahimahi, ‘ahi) were lower when only including the data from Carlisle et al. (2021) (Table S9). This additionally suggests an influence of sample size, and potentially a confounding effect of multiple prey occurring in similar isotopic space (Figure S7). Therefore, we can confidently infer that prey occurring in the defined epipelagic isotopic space make up a majority of false killer whales’ diet, but inferring prey-specific contributions within this space should be cautionary.

**Table S9.** Stable isotope mixing model results with four prey sources defined using prey data from only one study/source; posterior mean (95% credible interval; CrI) predicted proportion of each prey group to false killer whales' diet for the population and by social cluster.

| | Mean posterior proportion (95% CrI) | | | | |
|---|---|---|---|---|---|
| **Prey group**, *species* | Population | Cluster 1 | Cluster 2 | Cluster 3 | Cluster 4 |
| **Epipelagic predatory fish** | 0.577 (0.336-0.796) | 0.602 (0.336-0.835) | 0.542 (0.295-0.789) | 0.622 (0.340-0.856) | 0.593 (0.269-0.878) |
| *Mahimahi* | 0.057 (0.002-0.183) | 0.067 (0.001-0.261) | 0.055 (0.002-0.191) | 0.059 (0.001-0.214) | 0.044 (0.001-0.163) |
| *‘Ahi* | 0.080 (0.003-0.233) | 0.088 (0.002-0.285) | 0.088 (0.002-0.299) | 0.075 (0.002-0.252) | 0.063 (0.001-0.238) |
| *Ono* | 0.058 (0.002-0.186) | 0.063 (0.002-0.219) | 0.063 (0.002-0.241) | 0.057 (0.002-0.202) | 0.045 (0.001-0.178) |
| *A‘u* | 0.045 (0.002-0.147) | 0.047 (0.001-0.165) | 0.039 (0.001-0.138) | 0.045 (0.001-0.166) | 0.037 (0.001-0.150) |
| *Aku* | 0.064 (0.002-0.197) | 0.067 (0.001-0.229) | 0.062 (0.001-0.223) | 0.062 (0.001-0.217) | 0.055 (0.001-0.236) |
| *Uku* | 0.174 (0.018-0.380) | 0.158 (0.012-0.337) | 0.137 (0.012-0.321) | 0.201 (0.013-0.429) | 0.278 (0.011-0.613) |
| *Kahala* | 0.098 (0.007-0.267) | 0.113 (0.005-0.357) | 0.098 (0.005-0.328) | 0.123 (0.004-0.409) | 0.070 (0.003-0.226) |
| **Mesopelagic predatory fish** | 0.243 (0.104-0.410) | 0.236 (0.082-0.411) | 0.298 (0.110-0.481) | 0.211 (0.067-0.377) | 0.184 (0.058-0.342) |
| *‘Ahi palaha* | 0.067 (0.004-0.195) | 0.065 (0.002-0.209) | 0.089 (0.002-0.301) | 0.054 (0.002-0.178) | 0.054 (0.002-0.192) |
| *‘Ahi po‘onui* | 0.053 (0.002-0.173) | 0.051 (0.001-0.179) | 0.063 (0.001-0.236) | 0.045 (0.001-0.160) | 0.039 (0.001-0.141) |
| *A‘u ku* | 0.060 (0.002-0.194) | 0.060 (0.002-0.205) | 0.059 (0.002-0.210) | 0.062 (0.001-0.223) | 0.046 (0.001-0.173) |
| *Opah* | 0.063 (0.002-0.188) | 0.060 (0.001-0.199) | 0.087 (0.001-0.277) | 0.050 (0.001-0.163) | 0.046 (0.001-0.163) |
| **Reef-associated predatory fish** (*Ulua*) | 0.091 (0.007-0.218) | 0.066 (0.004-0.162) | 0.066 (0.003-0.162) | 0.086 (0.004-0.208) | 0.142 (0.003-0.323) |
| **Mesopelagic cephalopods** (*Purpleback flying squid*) | 0.089 (0.005-0.238) | 0.095 (0.003-0.260) | 0.095 (0.003-0.258) | 0.082 (0.003-0.231) | 0.081 (0.002-0.279) |

Mixing model outputs with the three-prey source model—where kahala and uku were grouped with ulua as reef-associated predatory fish—largely showed similar findings as the four-prey source model, where epipelagic predatory fish contributed the most to false killer whales' diet followed by mesopelagic prey and reef-associated predatory fish (Table S10). The movement of kahala and uku from the epipelagic to reef-associated predatory fish group naturally resulted in slightly higher proportions of reef-associated predatory fish in the three-prey source model (population posterior mean = 0.24, 95% CrI = 0.07 to 0.46) compared to the four-prey source model (population posterior mean = 0.05, 95% CrI = 0.00 to 0.15). This also resulted in lower proportions of epipelagic predatory fish across all social clusters (Table S10). Despite these differences, cluster-specific trends in dietary proportions were largely preserved: Clusters 1 and 2 had the highest proportions of mesopelagic prey, and slightly lower proportions of reef-associated predatory fish (Table S10). Similar to the four-prey source model, Cluster 4 had the lowest proportion of mesopelagic prey sources and highest proportion of epipelagic predatory fish in the three prey-source model (Table S10).

**Table S10.** Stable isotope mixing model results with three prey sources defined (epipelagic predatory fish = a‘u, ono, mahimahi, ‘ahi, aku; mesopelagic prey = a‘u ku, ‘ahi po‘onui, opah, ‘ahi palaha, purpleback flying squid; reef-associated predatory fish = kahala, uku, ulua); posterior mean (95% credible interval; CrI) predicted proportion of each prey group to false killer whales' diet for the population and by social cluster. Prey-specific contributions to diet are provided in Table S9 but note the potential confounding effect of multiple prey occurring in similar isotopic space (Figure S7).

| | **Posterior mean proportion of diet (95% CrI)** |
|---|---|

| | Epipelagic predatory fish | Mesopelagic prey | Reef-associated predatory fish |
|---|---|---|---|
| Population | 0.42 (0.17-0.66) | 0.34 (0.18-0.52) | 0.24 (0.07-0.50) |
| Cluster 1 | 0.41 (0.14-0.72) | 0.37 (0.17-0.58) | 0.22 (0.04-0.46) |
| Cluster 2 | 0.38 (0.13-0.66) | 0.43 (0.24-0.63) | 0.19 (0.03-0.41) |
| Cluster 3 | 0.43 (0.15-0.73) | 0.30 (0.11-0.51) | 0.28 (0.06-0.55) |
| Cluster 4 | 0.52 (0.10-0.85) | 0.23 (0.07-0.46) | 0.25 (0.03-0.65) |

Supporting Information References

Abecassis, M., Dewar, H., Hawn, D. & Polovina, J. (2012). Modeling swordfish daytime vertical habitat in the North Pacific Ocean from pop-up archival tags. *Mar. Ecol. Prog. Ser.*, 452, 219–236.

Arostegui, M.C., Gaube, P., Bowman, M., Nakamaru, K. & Braun, C.D. (2024). Fishery-independent and -dependent movement data aid in defining the stock structure of a data-deficient billfish. *Fish. Res.*, 271, 106923.

Arostegui, M.C., Gaube, P. & Braun, C.D. (2019). Movement ecology and stenothermy of satellite-tagged shortbill spearfish (*Tetrapturus angustirostris*). *Fish. Res.*, 215, 21–26.

Asher, J., Williams, I.D. & Harvey, E.S. (2017). An assessment of mobile predator populations along shallow and mesophotic depth gradients in the Hawaiian Archipelago. *Sci. Rep.*, 7, 3905.

Baird, R.W., Anderson, D.B., Kratofil, M.A. & Webster, D.L. (2021). Bringing the right fishermen to the table: Indices of overlap between endangered false killer whales and nearshore fisheries in Hawai‘i. *Biol. Conserv.*, 255, 108975.

Baird, R.W., Gorgone, A.M., McSweeney, D.J., Webster, D.L., Salden, D.R., Deakos, M.H., *et al.* (2008). False killer whales (*Pseudorca crassidens*) around the main Hawaiian Islands:

Long-term site fidelity, inter-island movements, and association patterns. *Mar. Mammal Sci.*, 24, 591–612.

Blum, J.D., Popp, B.N., Drazen, J.C., Anela Choy, C. & Johnson, M.W. (2013). Methylmercury production below the mixed layer in the North Pacific Ocean. *Nat. Geosci.*, 6, 879–884.

Brill, R.W., Block, B.A., Boggs, C.H., Bigelow, K.A., Freund, E.V. & Marcinek, D.J. (1999). Horizontal movements and depth distribution of large adult yellowfin tuna (*Thunnus albacares*) near the Hawaiian Islands, recorded using ultrasonic telemetry: implications for the physiological ecology of pelagic fishes. *Mar. Biol.*, 133, 395–408.

Bürkner, P.-C. (2018). Advanced Bayesian multilevel modeling with the R package brms. *The R Journal*, 10, 395–411.

Caputo, M., Kiszka, J.J., Andrianarivelo, N., Jonas, A., Andrianantenaina, B., Paz, V., *et al.* (2025). Trophic ecology of threatened sympatric coastal dolphins and other odontocetes in North-Western Madagascar. *Mar. Mammal Sci.*, e70027.

Carlisle, A.B., Allan, E.A., Kim, S.L., Meyer, L., Port, J., Scherrer, S., *et al.* (2021). Integrating multiple chemical tracers to elucidate the diet and habitat of cookiecutter sharks. *Sci. Rep.*, 11, 11809.

Carlisle, A.B., Goldman, K.J., Litvin, S.Y., Madigan, D.J., Bigman, J.S., Swithenbank, A.M., *et al.* (2015). Stable isotope analysis of vertebrae reveals ontogenetic changes in habitat in an endothermic pelagic shark. *Proc. R. Soc. B.*, 282, 20141446.

Choy, C.A., Popp, B.N., Hannides, C.C.S. & Drazen, J.C. (2015). Trophic structure and food resources of epipelagic and mesopelagic fishes in the North Pacific Subtropical Gyre ecosystem inferred from nitrogen isotopic compositions. *Limnol. Oceanogr.*, 60, 1156–1171.

Clark, C.T., Cape, M.R., Shapley, M.D., Mueter, F.J., Finney, B.P. & Misarti, N. (2021). SuessR: Regional corrections for the effects of anthropogenic $CO_2$ on $\delta^{13}$ C data from marine organisms. *Methods Ecol. Evol.*, 12, 1508–1520.

DeNiro, M.J. & Epstein, S. (1977). Mechanism of carbon isotope fractionation associated with lipid synthesis. *Science*, 197, 261–263.

Dewar, H., Prince, E.D., Musyl, M.K., Brill, R.W., Sepulveda, C., Luo, J., *et al.* (2011). Movements and behaviors of swordfish in the Atlantic and Pacific Oceans examined using pop-up satellite archival tags. *Fish. Oceanogr.*, 20, 219–241.

Domokos, R., Seki, M.P., Polovina, J.J. & Hawn, D.R. (2007). Oceanographic investigation of the American Samoa albacore (*Thunnus alalunga*) habitat and longline fishing grounds. *Fish. Oceanogr.*, 16, 555–572.

Dujon, A.M., Lindstrom, R.T. & Hays, G.C. (2014). The accuracy of Fastloc-GPS locations and implications for animal tracking. *Methods Ecol. Evol.*, 5, 1162–1169.

*FishBase*. (2025). . Available at: https://www.fishbase.se/. Last accessed 22 June 2025.

Flament, P. (1996). *Ocean Atlas of Hawai'i. Pacific Islands Ocean Observing System (PacIOOS)*. Available at: https://www.pacioos.hawaii.edu/education/ocean-atlas/. Last accessed 22 June 2025.

Fleming, C.H. & Calabrese, J.M. (2023). ctmm: continuous-time movement modeling.

Giménez, J., Ramírez, F., Almunia, J., G. Forero, M. & De Stephanis, R. (2016). From the pool to the sea: Applicable isotope turnover rates and diet to skin discrimination factors for bottlenose dolphins (*Tursiops truncatus*). *J. Exp. Mar. Biol. Ecol.*, 475, 54–61.

Gloeckler, K., Choy, C.A., Hannides, C.C.S., Close, H.G., Goetze, E., Popp, B.N., *et al.* (2018). Stable isotope analysis of micronekton around Hawaii reveals suspended particles are an important nutritional source in the lower mesopelagic and upper bathypelagic zones. *Limnol. Oceanogr.*, 63, 1168–1180.

Graham, B.S. (2007). Trophic dynamics and movements of tuna in the tropical Pacific Ocean inferred from stable isotope analyses. University of Hawaii at Manoa.

Jereb, P. & Roper, C.F.E. (2010). *Cephalopods of the world: An annotated and illustrated catalogue of cephalopod species known to date. Volume 2. Myopsid and Oegopsid Squids*. FAO Species Catalogue for Fishery Purposes. FAO, Rome, Italy.

Johnson, D.S. & London, J.M. (2025). ctmmUtils: auxillary functions for using the ctmm package efficiently.

Kratofil, M.A., Ylitalo, G.M., Mahaffy, S.D., West, K.L. & Baird, R.W. (2020). Life history and social structure as drivers of persistent organic pollutant levels and stable isotopes in Hawaiian false killer whales (*Pseudorca crassidens*). *Sci. Total Environ.*, 733, 138880.

Lam, C.H., Tam, C., Kobayashi, D.R. & Lutcavage, M.E. (2020). Complex dispersal of adult yellowfin tuna from the main Hawaiian Islands. *Front. Mar. Sci.*, 7, 138.

London, J.M. (2020). pathroutr: an R package for (re-)routing paths around barriers.

Lysy, M., Stasko, A.D. & Swanson, H.K. (2023). nicheROVER: niche region and niche overlap metrics for multidimensional ecological niches.

McGowen, M.R., Tsagkogeorga, G., Álvarez-Carretero, S., dos Reis, M., Struebig, M., Deaville, R., *et al.* (2020). Phylogenomic resolution of the cetacean tree of life using target sequence capture. *Syst. Biol.*, 69, 479–501.

Musyl, M.K., Brill, R.W., Boggs, C.H., Curran, D.S., Kazama, T.K. & Seki, M.P. (2003). Vertical movements of bigeye tuna (*Thunnus obesus*) associated with islands, buoys, and seamounts near the main Hawaiian Islands from archival tagging data. *Fish. Oceanogr.*, 12, 152–169.

Phillips, D.L., Inger, R., Bearhop, S., Jackson, A.L., Moore, J.W., Parnell, A.C., *et al.* (2014). Best practices for use of stable isotope mixing models in food-web studies. *Can. J. Zool.*, 92, 823–835.

Phillips, D.L., Newsome, S.D. & Gregg, J.W. (2005). Combining sources in stable isotope mixing models: alternative methods. *Oecologia*, 144, 520–527.

Polovina, J.J., Hawn, D. & Abecassis, M. (2008). Vertical movement and habitat of opah (*Lampris guttatus*) in the central North Pacific recorded with pop-up archival tags. *Mar. Biol.*, 153, 257–267.

Sackett, D.K., Drazen, J.C., Choy, C.A., Popp, B. & Pitz, G.L. (2015). Mercury sources and trophic ecology for Hawaiian bottomfish. *Environ. Sci. Technol.*, 49, 6909–6918.

Sackett, D.K., Drazen, J.C., Popp, B.N., Choy, C.A., Blum, J.D. & Johnson, M.W. (2017). Carbon, nitrogen, and mercury isotope evidence for the biogeochemical history of mercury in Hawaiian marine bottomfish. *Environ. Sci. Technol.*, 51, 13976–13984.

Schaefer, K.M. & Fuller, D.W. (2007). Vertical movement patterns of skipjack tuna (*Katsuwonus pelamis*) in the eastern equatorial Pacific Ocean, as revealed with archival tags. *Fish. Bull.*, 105, 379–389.

Sepulveda, C.A., Aalbers, S.A., Ortega-Garcia, S., Wegner, N.C. & Bernal, D. (2011). Depth distribution and temperature preferences of wahoo (*Acanthocybium solandri*) off Baja California Sur, Mexico. *Mar. Biol.*, 158, 917–926.

Smith, J.A., Mazumder, D., Suthers, I.M. & Taylor, M.D. (2013). To fit or not to fit: evaluating stable isotope mixing models using simulated mixing polygons. *Methods Ecol. Evol.*, 4, 612–618.

Stock, B.C., Jackson, A.L., Ward, E.J., Parnell, A.C., Phillips, D.L. & Semmens, B.X. (2018). Analyzing mixing systems using a new generation of Bayesian tracer mixing models. *PeerJ*, 6, e5096.

Stock, B.C. & Semmens, B.X. (2016). MixSIAR GUI User Manual.

Swanson, H.K., Lysy, M., Power, M., Stasko, A.D., Johnson, J.D. & Reist, J.D. (2015). A new probabilistic method for quantifying n-dimensional ecological niches and niche overlap. *Ecology*, 96, 318–324.

Tanaka, K.R., Schmidt, A.L., Kindinger, T.L., Whitney, J.L. & Samson, J.C. (2022). *Spatiotemporal assessment of* Aprion virescens *density in shallow main Hawaiian Islands waters, 2010-2019* (NOAA Technical Memorandum No. NMFS-PIFSC-132). U.S. Department of Commerce.

Velasquez-Vacca, A., Seminoff, J.A., Jones, T.T., Balazs, G.H. & Cardona, L. (2024). Trophic history of Hawaiian green turtles as revealed by stable isotope ratios ($\delta^{13}C$, $\delta^{15}N$ and $\delta^{34}S$) in the bones of museum specimens. *Aquat. Conserv.*, 34, e4063.

Vincent, C., McConnell, B.J., Ridoux, V. & Fedak, M.A. (2002). Assessment of Argos location accuracy from satellite tags deployed on captive gray seals. *Mar. Mammal Sci.*, 18, 156–166.

Whitney, N., Taquet, M., Brill, R.W., Girard, C., Schwieterman, G.D., Dagorn, L., *et al.* (2016). Swimming depth of dolphinfish (*Coryphaena hippurus*) associated and unassociated with fish aggregating devices. *Fish. Bull.*, 114, 426–434.

Ylitalo, G.M., Baird, R.W., Yanagida, G.K., Webster, D.L., Chivers, S.J., Bolton, J.L., *et al.* (2009). High levels of persistent organic pollutants measured in blubber of island-associated false killer whales (*Pseudorca crassidens*) around the main Hawaiian Islands. *Marine Pollution Bulletin*, 58, 1932–1937.

Zaeschmar, J.R. & Baird, R.W. (2025). False killer whale *Pseudorca crassidens* (Owen, 1846). In: *Ridgway and Harrison's Handbook of Marine Mammals* (ed. Jefferson, T.). Elsevier.
